# Convolutional networks for supervised mining of molecular patterns within cellular context

**DOI:** 10.1101/2022.04.12.488077

**Authors:** Irene de Teresa, Sara K. Goetz, Alexander Mattausch, Frosina Stojanovska, Christian E. Zimmerli, Mauricio Toro-Nahuelpan, Dorothy W.C. Cheng, Fergus Tollervey, Constantin Pape, Martin Beck, Anna Kreshuk, Julia Mahamid, Judith Zaugg

**Affiliations:** Structural and Computational Biology Unit, European Molecular Biology Laboratory, Heidelberg, Germany.; Computer Science and Artificial Intelligence Lab, ENGIE Lab Crigen, Stains, France.; Collaboration for Joint PhD Degree between EMBL and Heidelberg University, Faculty of Biosciences, Heidelberg, Germany.; Institute of Pharmacy and Molecular Biotechnology, Heidelberg University, Heidelberg, Germany.; Max Planck Institute of Biophysics, Department of Molecular Sociology, Frankfurt, Germany.; Santiago GmbH & Co. KG, Willich, Germany.; Cell Biology and Biophysics Unit, European Molecular Biology Laboratory, Heidelberg, Germany.; Genome Biology Unit, European Molecular Biology Laboratory, Heidelberg, Germany.; Institute for Computer Science, Universität Göttingen

## Abstract

Cryo-electron tomograms capture a wealth of structural information on the molecular constituents of cells and tissues. We present DeePiCt (Deep Picker in Context), an open-source deep-learning framework for supervised structure segmentation and macromolecular complex localization in cellular cryo-electron tomography. To train and benchmark DeePiCt on experimental data, we comprehensively annotated 20 tomograms of *Schizosaccharomyces pombe* for ribosomes, fatty acid synthases, membranes, nuclear pore complexes, organelles and cytosol. By comparing our method to state-of-the-art approaches on this dataset, we show its unique ability to identify low-abundance and low-density complexes. We use DeePiCt to study compositionally-distinct subpopulations of cellular ribosomes, with emphasis on their contextual association with mitochondria and the endoplasmic reticulum. Finally, by applying pre-trained networks to a HeLa cell dataset, we demonstrate that DeePiCt achieves high-quality predictions in unseen datasets from different biological species in a matter of minutes. The comprehensively annotated experimental data and pre-trained networks are provided for immediate exploitation by the community.

## Introduction

Cryo-Electron Tomography (cryo-ET) produces three-dimensional (3D) snapshots of cellular landscapes at molecular resolution, making it possible to investigate the structural and functional states of macromolecular complexes in their native environment, and to unveil how different macromolecular populations interact with other cellular structures^1–8^. With improved instrumentation, sample preparation protocols, and automation, high-quality in-cell cryo-ET data are rapidly being generated, opening the possibility to conduct high-throughput studies^9–12^. However, due to the complex and crowded nature of the intracellular milieu, together with limitations arising from cryo-ET image acquisition (low signal-to-noise ratio (SNR) and incomplete angular sampling) data mining of 3D cryo-ET volumes remains a major bottleneck^13, 14^. Data mining in terms of reliable identification of a relatively homogenous set of macromolecular complexes constitutes a fundamental prerequisite for structural analysis^13–15^.

A range of available semi-automated methods for segmentation of cellular structures and localization of macromolecular complexes (from here on, particles) in cryo-ET datasets are broadly classified as template-based and template-free approaches^16^. Traditional Template Matching (TM)^17^ is a commonly applied computational approach and is based on a point-wise numerical computation of a similarity coefficient (cross-correlation) to a known template of the complex in question. TM is accurate in the localization of ribosomes and cytoskeletal filaments, but fails at identifying smaller or less dense particles^18^, and is computationally intensive. On the other hand, current template-free methods implementing classical image processing approaches are purposely designed for and thus limited to specific molecular shapes, or to special configurations in which particles are associated with large cellular structures such as membranes or microtubules^16, 19, 20^. These methods together with TM require manual inspection and are therefore laborious and time-consuming. The advent of deep learning methods, and particularly convolutional neural networks (CNNs)^21–23^, has enabled developing more generally applicable and automated approaches to accelerate the tasks of segmentation and localization in cryo-ET. The first of such methods was a two-dimensional (2D) CNN for semantic segmentation of ribosomes and large structures, such as cellular organelles or membranes^24^. However, its 2D nature makes it less suitable for particle localization, where probing the 3D structure becomes beneficial. More recently, DeepFinder, a fully supervised method based on the U-Net architecture^25^ for multi-class semantic segmentation, has positioned itself as the state-of-the-art in automated particle localization in both simulated and real cryo-ET datasets^26–29^. Remaining limitations include the identification of less prevalent particles, as well as the interpretation of the obtained predictions within their cellular context. Furthermore, the field lacks publicly available non-synthetic expert-annotated cryo-ET data to train new models and to benchmark the available methods.

Here we present DeePiCt (Deep Picker in Context), our open-source software which synergizes supervised convolutional networks for segmentation of cellular compartments (organelles or cytosol) and structures (membranes or cytoskeletal filaments), and localization of particles. We generated a set of comprehensively annotated tomograms of wild-type *S. pombe* for training and benchmarking our method, which we make publicly available to overcome the critical limitations arising from the absence of publicly available annotated experimental datasets. From here on, we refer to the term *network* as the deep learning algorithm itself and *model* as the algorithm once already trained. We provide DeePiCt models, trained on this dataset which show high data-mining performance and can be readily applied across species and datasets.

## Results

### DeePiCt: A deep-learning approach for automated compartment segmentation and particle localization

DeePiCt is based on deep learning technologies. It combines a 2D CNN for segmentation of cellular compartments and a 3D CNN for particle localization, membrane, and cytoskeletal filament annotations (**Fig. 1**). This synergy allows for more precise particle picking and interpretation of their cellular context. The 2D and 3D CNN are adapted from the original U-Net architecture^25^ (**Fig. 1a, Supplementary Note 1**). Since cellular compartments are easily recognizable in 2D image data, the compartment segmentation network operates in 2D, which has the advantage of requiring little training data. The 2D CNN employs a fixed depth (D) of 5 (4 max-pooling layers) and 16 initial filters (IF) (**Fig. 1a**). Our implementation of the 3D CNN allows multi-label learning (**Supplementary Note 1**) and adjustable architectural parameters D, number of IF, a batch normalization layer (BN), and the dropout parameter in the encoder and decoder paths (ED and DD, respectively) (**Fig. 1a**). These parameters can be set according to the amount and quality of training data, and the shape complexity, size and abundance of the particle of interest as suggested in **Supplementary Table 1**. In general, larger particles benefit from larger values of D to increase the network’s receptive field, and particles with a low-density print (low SNR with respect to the surrounding context) require larger IF values. The remaining optional layers, BN and dropout, are well known techniques in computer vision to avoid overfitting^30, 31^.

**Figure 1.**
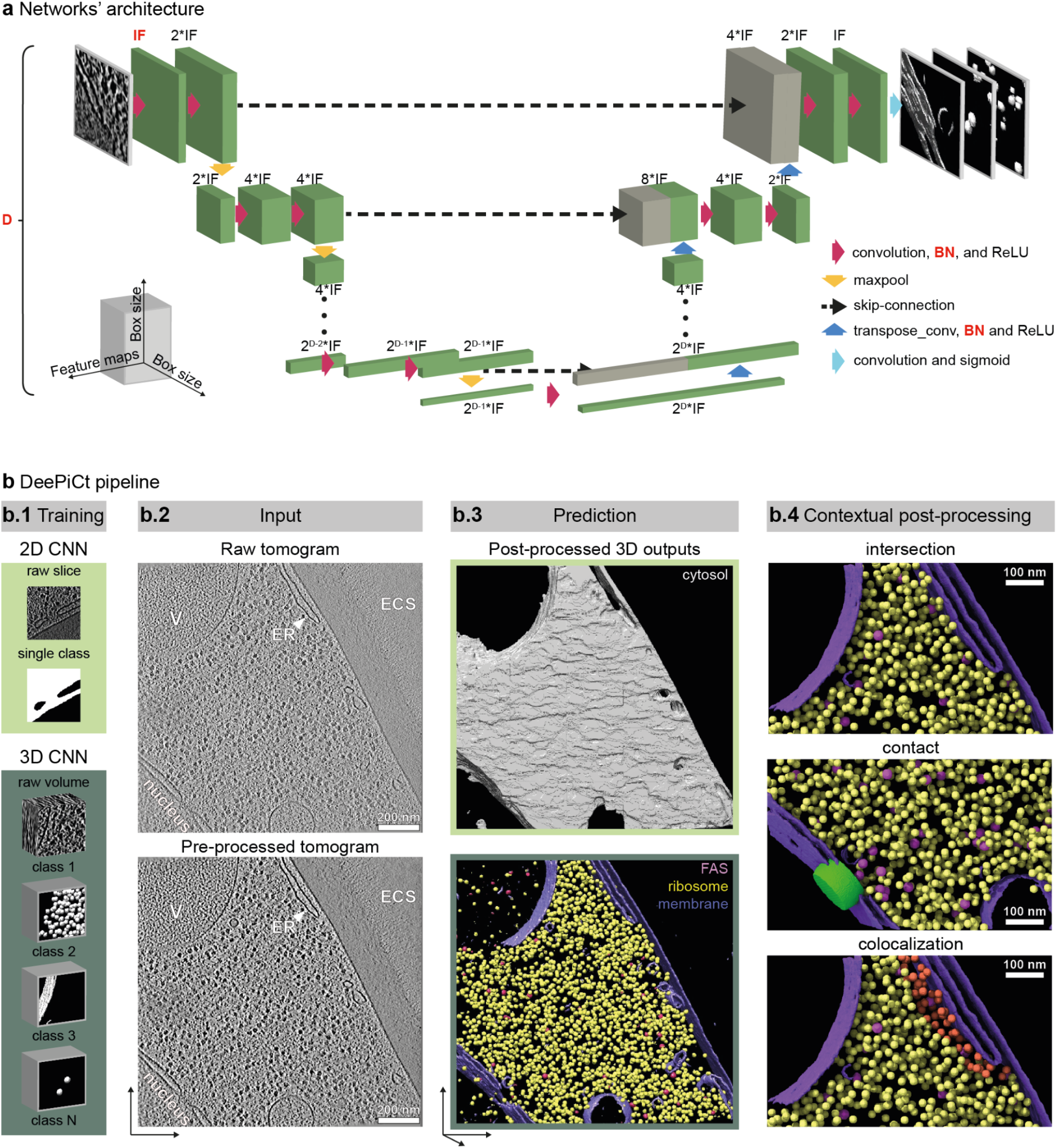
DeePiCt 2D and 3D CNN architecture implemented in an automated workflow combining compartment segmentations and particle localizations in cryo-ET data. **a**. The CNN U-Net architecture: the 2D network performs all tensor operations on the two-dimensional spatial coordinates, with D=5 and IF=16; the 3D network performs tensor operations on the three spatial dimensions, and the architectural hyperparameters in red can be set by the user. **b**. The DeePiCt pipeline is used to train and predict various structural features in cryo-electron tomograms. **b.1** The DeePiCt pipeline consists of two independent CNNs: a 2D network for compartment segmentation and a 3D network for particle localization. **b.2** Trained networks are applied to input tomograms which can be pre-processed with a spectrum-matching filter to improve image contrast. The example 2D tomographic slice visualizes the cytoplasm with the endoplasmic reticulum (ER), vacuoles (V), nucleus and extracellular space (ECS). **b.3** DeePict raw predictions for cytosol, membranes, ribosomes and fatty acid synthase (FAS) are post-processed by thresholding, cluster size and centroid fitting. **b.4** The outputs of the two networks can be combined to include the cellular context by intersecting particle predictions with cytosol masks (top), selecting particles (NPC, green) in contact with specific membranes (nuclear envelope, middle), and to identify particles (e.g. ribosomes, orange) associated with specific organelles (e.g. ER).

For training, each network requires tomograms and corresponding 3D binary masks, which represent the segmentation of the structures of interest, e.g. organelle segmentations for the 2D U-Net and spheres representing particles for the 3D U-Net (**Fig. 1b.1)**. The raw input tomograms are optionally preprocessed using an amplitude spectrum equalization filter (**Fig. 1b.2, Extended Data Fig. 1**) to enhance image contrast by matching the amplitude spectrum of the input tomogram to a preselected target amplitude spectrum distribution (**Extended Data Fig. 1**). Both CNNs use the Adam optimizer algorithm and the Dice Loss function for training (**Supplementary Note 1**). In the 2D network, tiles are randomly flipped and rotated in 90-degree increments during training to improve generalization. For the 3D CNN, we implemented a data augmentation strategy by applying a number of optional random transformations to the input images (**Supplementary Note 1**). In our experience, more than 700 annotated instances are required for particle learning in the 3D CNNs, while about 5 tomograms are sufficient for membrane segmentation. In the case of 2D CNNs, 6 fully segmented tomograms are enough for training networks identifying either a cytosol class or a single class including all cellular organelles.

For predicting, the trained networks receive as input unseen tomograms pre-processed in the same way as the training data, and output 3D probability maps that are subsequently post-processed (**Fig. 1b.3**). Automated post-processing for the 2D network includes the combination of predicted 2D slices into a 3D volume, application of a 1D Gaussian filter along the z-axis for smoothing the predictions across the z slices, and thresholding (user-definable, default=0.75) (**Extended Data Fig. 2a-c**). The 3D CNN post-processing entails the generation of a predicted segmentation. To that end, the 3D CNN output is thresholded by a probability value set by the user and then clustered. The clustered output can be integrated with contextual information from a binary map representing a tomographic region (e.g. the cytosol segmentation from the 2D CNN). The mode of integration can be chosen among three different options: *intersection*, *contact*, or *colocalization*, depending on the users’ specific application (**Fig. 1b.4**, **Supplementary Note 1, Supplementary Fig. 1**). In the case of particle localization, the clusters’ centroids are used to obtain a list of the particle coordinates; their orientations and structural features can then be obtained by subtomogram analysis in external softwares (e. g. Warp^32^, M^33^, RELION^34^, Dynamo^35^, PyTom^36^ or EMAN2^37^). For more details on the method, we refer to **Supplementary Note 1** and our github repository (Code and data availability).

### Generation of comprehensive ground-truth annotations in *S. pombe*

The ultimate requirement for developing and benchmarking any deep neural network is the existence of a ground truth (gt) non-synthetic dataset. Since this was not available at the outset of this study, we first created a comprehensive annotation of cellular features in wild-type *S. pombe* representing diverse structures, sizes and abundances.

In cryo-ET data, the complete annotation of a given cellular structure or particle species is a challenging process, even for large compartments or complexes with a clear structural signature like the ribosome. To achieve the most comprehensive annotation possible, we devised an iterative workflow to mine patterns in three different datasets (**Online Methods**). In a set of 10 tomograms acquired by combining defocus and Volta potential phase plate (VPP^38, 39^) and a second set of 10 tomograms using defocus only (DEF), we annotated ribosomes (RIBO), fatty acid synthases (FAS), membranes, organelles and cytoplasm (**Fig. 2, Extended Data Tables 1-3**). For the nuclear pore complex (NPC), given its structural flexibility, low abundance (on average 3 per tomogram), and large size, a third dataset denoted by DEF* was used^40^. It consists of 127 tomograms featuring a total of 2,830 NPC subunits (∼354 NPCs) to ensure enough training data and is composed of two subsets: one of higher and one of lower quality classified based on lamella thickness and tilt-series alignment error. A detailed description of the ground truth construction is available in the **Online Methods**.

**Figure 2.**
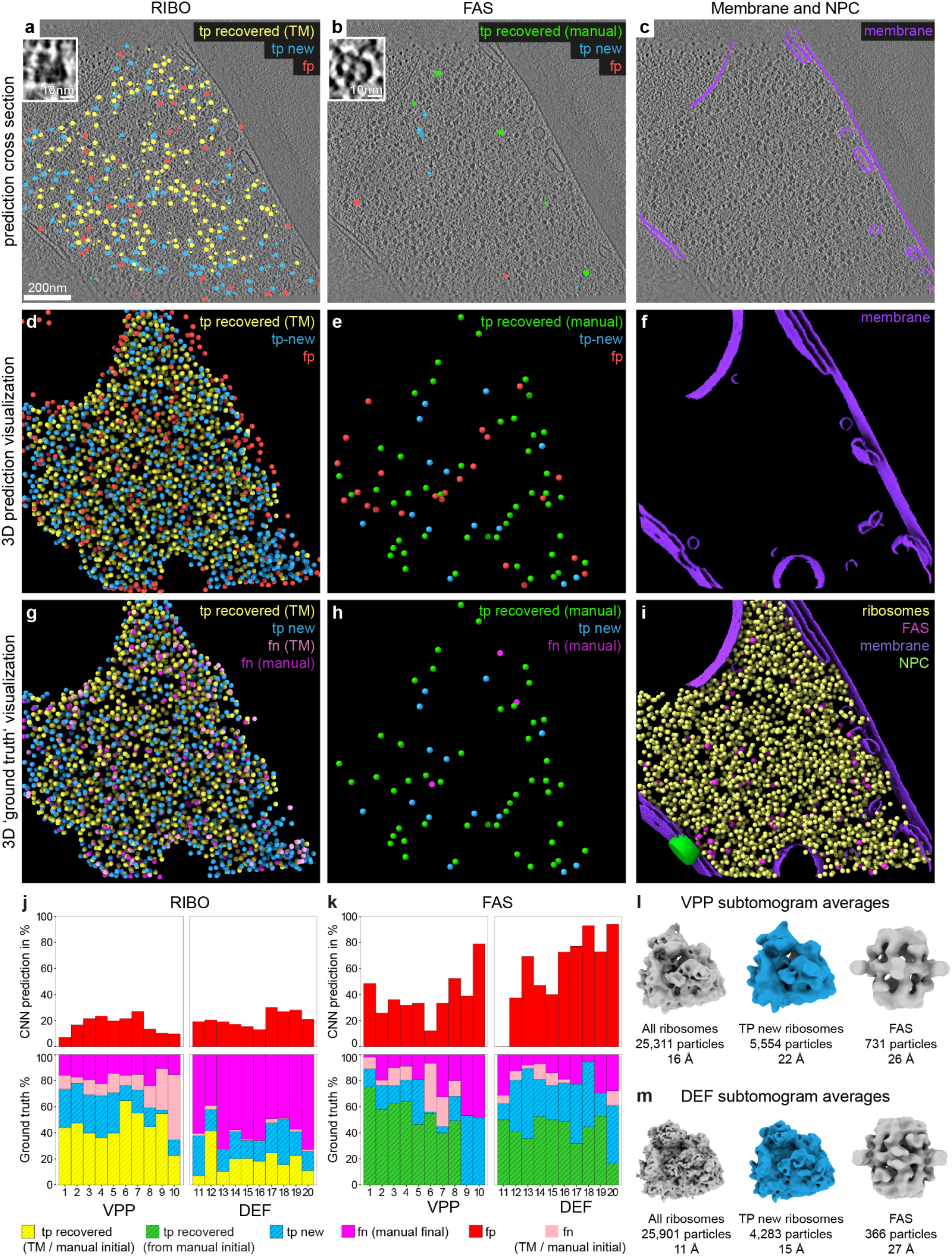
Iterative comprehensive annotation of ground truth for non-synthetic data. The three columns (left to right) show: the annotation process for RIBO (**a, d, g**), FAS (**b, e, h**), membranes (**c, f**). **a-f** Show the cumulative predictions of 3 rounds of DeePiCt for RIBO and one round for FAS as cross-section overlaid on a tomographic slice (**a-c**) and in 3D view (**d-f**). Particles are classified as true positives recovered from the initial annotation (tp recovered; yellow for RIBO; green for FAS), new true positives provided by DeePiCt predictions (tp new; blue), and the corresponding false positives (fp; red). **g, h**. 3D views of the resulting ground truth, including false negatives (fn) distinguishing unrecovered fn from the initial annotations (pink) and final round of manual picking (salmon). **i**. Combined ground truth annotation of RIBO, FAS, membranes, and NPC (green cylindrical mask, manual annotation). **j-k**. Relative contributions of the DeePiCt rounds for RIBO (**j**) and FAS (**k**) identification are plotted across 10 VPP and 10 DEF *S. pombe* tomograms: contribution of step 1 (yellow: TM and manual annotation for RIBO; green: manual annotation for FAS; salmon: initial annotations not detected by DeePiCt), step 2 (blue); and step 3 (pink bars). For FAS the initial annotation was not based on TM, and instead was purely manual. **l**, **m**. Subtomogram averages of all RIBO, exclusively DeePiCt-detected (tp new; blue) and all FAS particles in VPP and DEF ground truth, respectively.

The resulting ground truth RIBO annotation in VPP consisted of 25,311 particles. An initial TM annotation contributed 61.6% (1,559 particles per tomogram (ppt), on average) of the final ground truth, whereas iterative rounds of DeePiCt on the incomplete ground truth contributed to a total of additional 21.36% (541 ppt on average) and final manual annotations to 17.05% (431 ppt on average) (**Fig. 2j**, **Extended Data Fig. 3a,e, Extended Data Table 2**).

The final ground truth annotation of FAS in the 10 VPP datasets amounts to 731 particles, from which 58.96% (43 ppt on average) constituted the initial annotations from manual picking, as TM failed. The combined predictions of DeePiCt on the incomplete ground truth contributed to 21.88% (16 ppt on average). For FAS we performed only one round of DeePiCt using the initial manual annotations, since we observed a high false positive rate in subsequent rounds, which is likely a consequence of the natural lower abundance of FAS in the tomograms and its low imprint density. The final manual picking added a significant fraction of 19% (14 ppt on average) (**Extended Data Fig. 3c,e**, **Extended Data Table 3**).

The final ground truth RIBO annotations on the DEF dataset sum a total of 25,901 particles across the 10 tomograms, from which only 19.23% (498 ppt on average) were included in the initial TM annotation (step 1), 19.39% (502 ppt on average) were a contribution of DeePiCt on incomplete ground truth, and 61.36% (1,590 ppt on average) were manually identified (**Fig. 2k-m, Extended Data Fig. 3b,e**), highlighting the value of our carefully curated annotations. In the case of FAS, the ground truth annotations across the 10 tomograms amount to 366 particles, from which 48.63% (18 ppt on average) came from the initial manual annotations (as TM failed), 36.88% (14 ppt on average) from DeePiCt trained on incomplete ground truth, and 14.48% (5 ppt on average) from the final manual picking (**Fig. 2k-m**, **Extended Data Fig. 3d,f**). Overall, a final round of manual picking for RIBO ground truth annotation was notably more crucial in DEF compared to VPP.

Total numbers of RIBO in both acquisition types are comparable, while for FAS fewer particles were detected in DEF likely due to the lower SNR despite the application of an equalization filter (**Extended Data Fig. 3, Extended Data Table 2-3, Supplementary Note 2**). FAS annotations were similarly distributed among the ground truth construction steps in the two acquisition conditions (**Extended Data Fig. 3**). Comparison to previous proteomics studies^41, 42^ shows that our annotations of fully assembled FAS in cryo-electron tomograms is in agreement with levels of FAS-*α* and FAS-*β* quantified by means of mass spectrometry, while the abundance of cytosolic ribosomal proteins detected by mass spectrometry are underestimated (**Extended Data Fig. 4**).

To assess the quality of the RIBO and FAS particle annotations in the VPP and DEF datasets, we processed the tomograms in Warp and performed structural 3D classification and refinement of the subtomograms in RELION (**Extended Data Tables 4-5, Supplementary Note. 2, Supplementary Fig. 2-4**). Subtomogram averages of RIBO particles detected only by DeePiCt (’tp new’), represent 80S ribosomes, consistent with maps of all cytosolic RIBO particles for both VPP and DEF datasets (**Fig. 2l-m**, blue and gray, respectively). 3D classifications performed in both data types for FAS demonstrated that the 3D CNN together with manual annotations were capable of recovering these challenging shell-like structures independent of the data acquisition type (**Supplementary Note 2, Supplementary Fig. 3**).

In conclusion, using DeePiCt, we provide comprehensive annotations for large macromolecular complexes and cellular structures in *S. pombe* cryo-electron tomograms acquired with VPP and DEF, which set the ground for benchmarking the method’s performance and for future developments in particle detection approaches.

### Performance analysis and hyperparameter tuning of the DeePiCt workflow

#### Performance analysis of 2D CNN in VPP

For the 2D CNN, basic hyperparameter tuning had close to no effect and a fixed architecture produced sufficiently good segmentations. Hence, no extensive hyper-parameter tuning was performed. The 2D CNN’s performance was evaluated on 10 manually annotated ground-truth VPP tomograms in a 5-fold cross-validation (CV). For each CV fold, two tomograms were split off from the training data to evaluate model performance during training. Two binary segmentation tasks were evaluated: segmentation of all organelles, *i.e.* all membrane-enclosed organelles and the nucleoplasm, and segmentation of the cytosol. The results show that the 2D CNN achieves high areas under the precision-recall curve (AUPRC, **Supplementary Note 1**), with a median AUPRC of 0.98 for cytosol and 0.92 for organelles (**Fig. 3a**, **Extended Data** **Fig. 5**).

**Figure 3.**
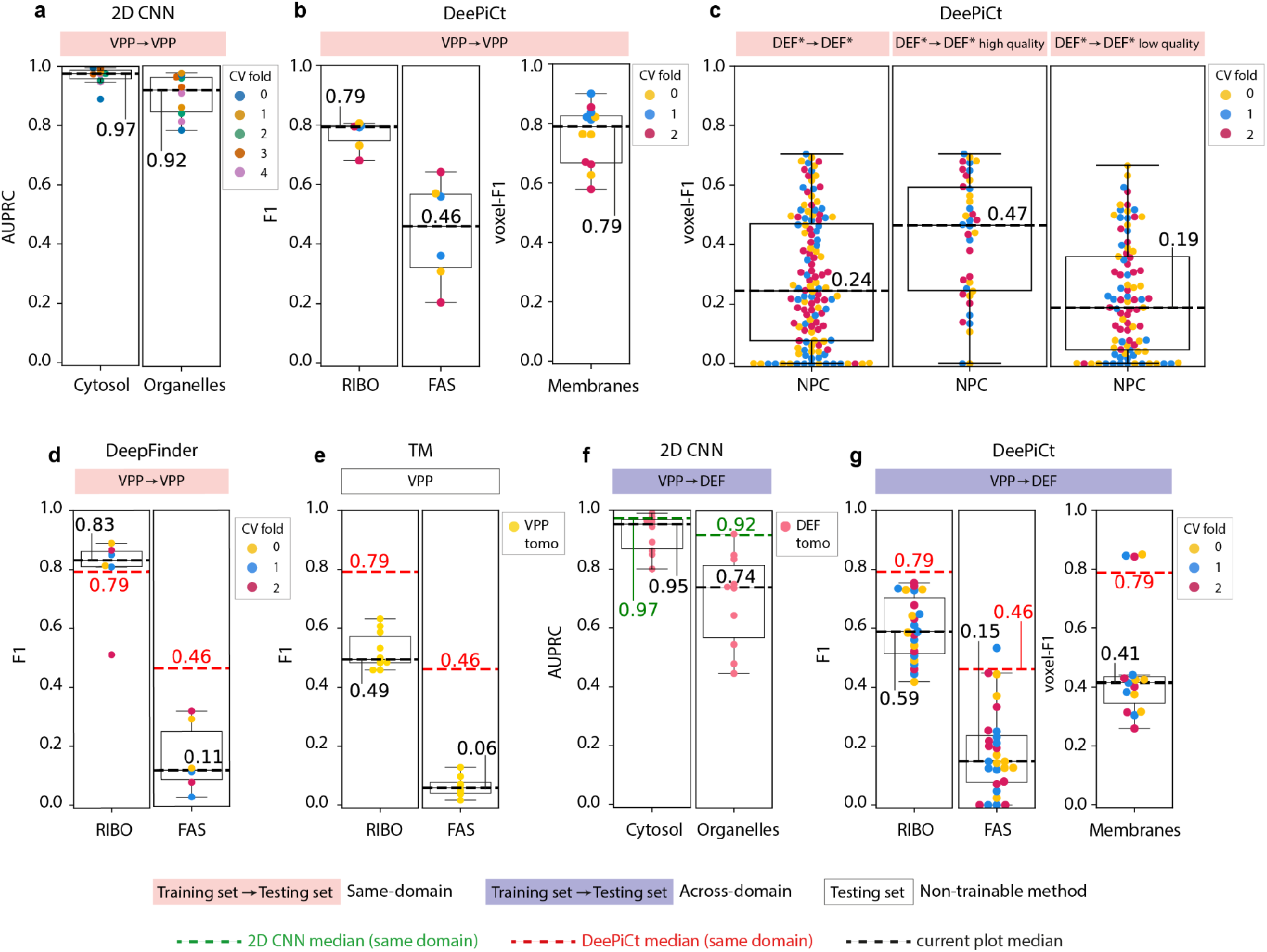
DeePiCt performance, cross-domain generalization and comparison with other methods. **a**. 2D CNN performance results for organelle and cytosol segmentation under a 5-fold CV scheme, when training and testing datasets are both in the same domain (VPP). The median AUPRC scores are indicated (black dotted lines). **b**. Performance of DeePiCt for the same-domain setting. The RIBO localization, FAS localization, and membrane segmentation tasks are shown (left to right), under a 3-fold CV scheme. In each case, the corresponding architectures of the 3D CNN were optimized by hyperparameter tuning (**Supplementary Table 1**, **Extended Data Fig. 6**). The median F1 score for RIBOS and FAS, and a median voxel-F1 for membranes are indicated (black dotted lines). **c**. Same-domain NPC segmentation results using a 3-fold CV scheme. From left to right: performance in all DEF* tomograms, high quality DEF* tomograms, and lower quality DEF* tomograms. **d**-**e**. DeepFinder (**d**) and TM (**e**) particle localization results for RIBO and FAS localization. The median F1 values are indicated in each case (black dashed line), along with corresponding values of DeePiCt’s performance for comparison (red dashed line). **d**. DeepFinder results are based on the same 3-fold CV scheme as in **b**, by training a multiclass DeepFinder network that simultaneously segments FAS and RIBO. **e**. TM results are shown for the VPP tomograms. **f**-**g**. Results of the cross-domain generalization for the 2D CNN (**f**) and DeePiCt (**g**) (training on 10 VPP tomograms, testing on 10 DEF tomograms). Dashed lines and their numerical labels indicate median performance values either of the corresponding plot (black), or the corresponding experiment performed in the same domain with DeePiCt (red), or in the same domain with the 2D CNN (green).

The 2D CNN showed an average false positive rate (FPR) of 0.054 for cytosol and 0.024 for organelle segmentation, with the post-processing step providing marginal improvements (reducing FPR to 0.049 for cytosol and 0.021 for organelles, **Extended Data Fig. 2**). In addition to the two segmentation tasks described here, the 2D CNN has the potential to segment individual organelle types when provided with sufficient amounts of training data (**Extended Data Table 1**). Operating in 2D rather than 3D results in only little memory requirement (<1GB at the default batch size) and short training time (∼15 minutes to train on ten tomograms using a Nvidia V100 GPU; **Online Methods**).

#### Hyperparameter tuning for the 3D CNN

The 3D CNN is built on a flexible architecture for which we performed hyperparameter tuning. We characterized the effect of varying the adjustable hyperparameters (**Online Methods**) on the DeePiCt workflow using a CV approach for RIBO, FAS and membranes in the VPP dataset, and for NPC in the DEF* dataset.

In each case we combined the 3D CNN segmentation of the target structure with an appropriate *region mask* (**Supplementary Fig. 1**, **Supplementary Table 1**) and determined the 3D CNN hyperparameter combination that optimizes the task of localization or segmentation (**Extended Data Fig. 6**, **Supplementary Table 1**). Our results show that the best hyperparameter selection for the 3D CNN is structure-dependent, and not only related to the size of the complex or to the receptive field of the network (a function of the depth parameter D), but also to the particle’s density, symmetry and abundance, among others. Therefore, the architecture flexibility of our implementation of the 3D CNN is essential for the accurate segmentation of a diverse set of biological structures. In particular, BN shortens the learning speed (and notably improves the performance results in the case of NPCs and FAS). Dropout layers and a number of data augmentation strategies (Gaussian noise, salt-and-pepper noise, rotations, elastic transformations, see **Supplementary Note 1**) did not improve performance.

#### Performance of DeePiCt in the same-domain setting

Once the 3D CNN architectures which optimized the tasks of particle localization (for FAS and RIBO) and structure segmentation (membranes and NPC) were determined, we analyzed the performance of the DeePiCt workflow in the same-domain setting (*i.e.* training and testing in the same domain). The 3-fold CV tests in VPP (**Fig. 3b, Extended Data Fig. 7a-c**), showed that it achieved a performance F1 score between 0.68-0.80 for RIBO (median F1 of 0.79), 0.21-0.64 for FAS (median F1 of 0.46) and 0.58-0.90 for membranes (median voxel-F1 of 0.71). The results in DEF* for NPC segmentation (**Fig. 3c**) show that, even if the network is trained on the whole DEF* dataset (23% of which is high quality data, **Online Methods**), the performance of the method is clearly different in the high-versus lower-quality DEF* tomograms: with a median voxel-F1 of 0.47 in high-quality data, while a median voxel-F1 of 0.19 in lower-quality DEF* tomograms.

### Comparison of DeePiCt to state-of-the-art particle localization tools

To benchmark DeePiCt in the particle localization task, we compared it to TM and DeepFinder in the tasks of RIBO and FAS in the VPP dataset (**Fig. 3d,e**). Following the suggestions by the authors^28^, DeepFinder was trained in a multi-class fashion, where both particles were learnt simultaneously by a single network. For the training, we used the same 3-fold CV splitting of the VPP datasets as for DeePiCt. In RIBO localization, DeepFinder achieved a median F1 of 0.83, comparable to DeePiCt. For FAS localization, DeepFinder achieved a median F1 of 0.11 (significantly below DeePiCt, t-test p-value of 0.007; **Fig. 3d**). It is worth mentioning that single-class DeepFinder networks tested for FAS localization completely failed by not localizing any particles. DeePiCt needs ∼17h for training vs ∼3h for DeepFinder, while predicting and post processing is equally fast for both (∼500 clusters/minute) (**Online Methods**). Furthermore, DeePiCt’s trained networks (models) for all structures mentioned in this work are open source and publicly available (Code and data availability).

Finally, the analysis of TM performance was done with the raw output with respect to the top 2000 peaks in the case of RIBO and to the top 1000 peaks in the case of FAS. TM also takes several hours (s. parameters in Online Methods) and our results confirm that TM has lower performance than both deep-learning methods for the localization of RIBO and FAS, completely failing at the latter (**Fig. 3e**).

### Domain generalization of DeePiCt across image acquisition conditions

The DEF data is characterized by a lower SNR and lower contrast in the image, as compared to VPP (**Supplementary Note 2**). This makes the task of particle localization and structure segmentation more challenging, but has the advantage that the resulting subtomogram averages generally provide better resolutions^43^. The comprehensive ground truth construction in DEF enabled us to study the generalization potential of the DeePiCt workflow across image acquisition conditions, from VPP to DEF data. To that end, we predicted cytosol, organelles, membranes, FAS and RIBO in 10 DEF tomograms, with networks trained in the yeast VPP data (**Fig. 3f-g**).

Compared to segmentation of VPP data, the 2D CNN shows slightly poorer performance on DEF data for cytosol (**Fig. 3a,f**). By employing amplitude spectrum equalization for the 2D CNN, the domain generalization capabilities greatly improved for both cytosol (from initial median AUPRC=0.82 to median AUPRC=0.95) and organelles (from AUPRC=0.42 to 0.74) (**Fig. 3f**, **Extended Data** **Fig. 5**). For DeePiCt, the performance scores in this domain generalization setting also showed a decay with respect to the same domain setting: with a median F1 of 0.59 for RIBO, a median F1 of 0.15 for FAS, and a median voxel-F1 of 0.41 for membranes (**Fig. 3g**). Importantly, as in the case of the 2D CNN, the spectrum equalization filter in the preprocessing step improved the domain generalization results (**Extended Data Fig. 7d-f**). Furthermore, we confirmed that using the cytosol predictions as a *region mask* to clean particle annotations also improved the performances (**Extended Data Fig. 7g-h**). An example of DeePict segmentation outcome in the cross-domain setting is shown in **Fig. 4a-b** and visually resembles the ground truth annotation (**Extended Data Fig. 8a**). The DeePiCt predictions and the ground truth annotations enabled us to analyze the DEF tomograms in the same way as we had done for the VPP data, by downstream 3D structural classification and refinement (**Fig. 5**).

**Figure 4.**
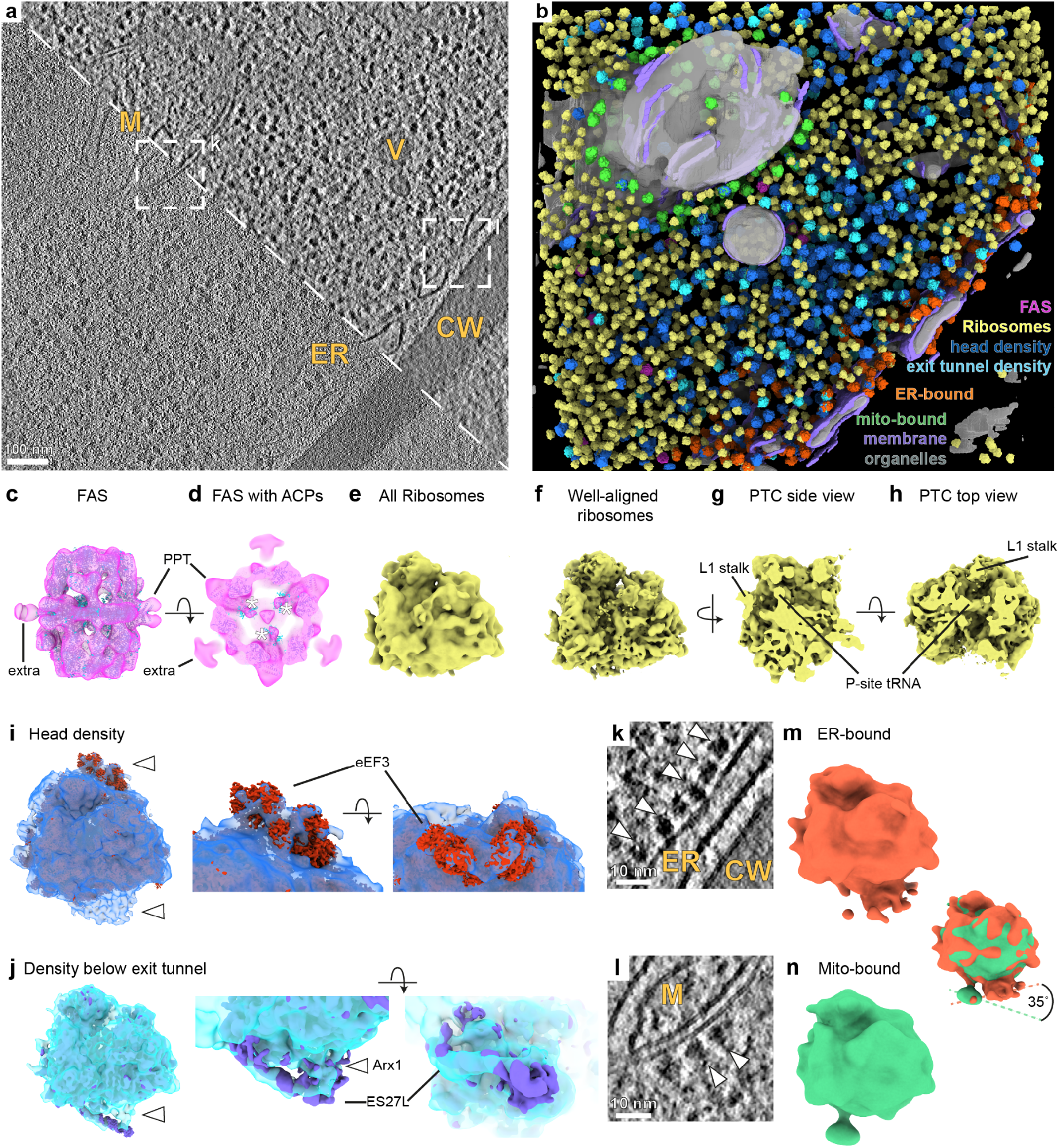
DeePiCt enables exploration of macromolecular complexes in their cellular context. **a**. 2D slice of a DEF *S. pombe* cytoplasm tomogram. Mitochondrion (M), vesicle (V), the ER and the cell wall (CW) are detectable in the raw (bottom left) and with improved contrast after amplitude spectrum equalization (top right). **b**. DeePiCt predictions generated with models trained in VPP data. Organelles (gray), membranes (purple), FAS (pink, **c**, **d**), ribosomes (yellow, **e**-**h**) and subsets classified in RELION (head density, dark blue, **i;** exit tunnel density, bright blue, **j)**, and within 25 nm to mitochondria (mito-bound, green, **m**) and the ER (ER-bound, orange, **n**). **c**. FAS subtomogram average (pink) fits the *S. cerevisiae* structure (cyan) including PPT domains. An extra density cannot be assigned. **d**. Cross-section of **c** close to the alpha-wheel with three densities fitting ACPs (white asterisks). **e**. Subtomogram average of all ribosomes from 10 DEF tomograms. **f**. Well-aligned ribosome subset detected by hierarchical 3D classification in RELION and refined in M. **g, h**. Slices through (**f**) reveal the PTC with a P-site tRNA and the L1 stalk facing the E-site. **i**. Ribosome subclass with additional densities (white arrowheads) close to the head of the small ribosomal subunit which fits eEF3 (red), and close to the ribosomal exit tunnel. **j**. Ribosomes classified for a density below the ribosomal exit tunnel (white arrowhead). The *S. cerevisiae* ribosome with ES27L in a particular configuration (left, purple) connects to the additional exit-tunnel density which fits Arx1 bound to the 60S pre-ribosome (middle and right, purple). **k**, **l**. Different z slices of the tomogram in **a** (white dashed boxes) show ribosomes bound to the ER (**k**) and a mitochondrion (**l**). **m**. ER-bound ribosomes from 7 tomograms show a connection of the peptide exit tunnel of the large subunit to the membrane density. **n**. Mitochondria-bound ribosomes from 3 tomograms show a linker connecting the large subunit, at a site close to the small subunit, to the membrane density. Overlay of the ribosomes in **m** and **n** shows different interfaces with the respective organelle membranes.

**Figure 5.**
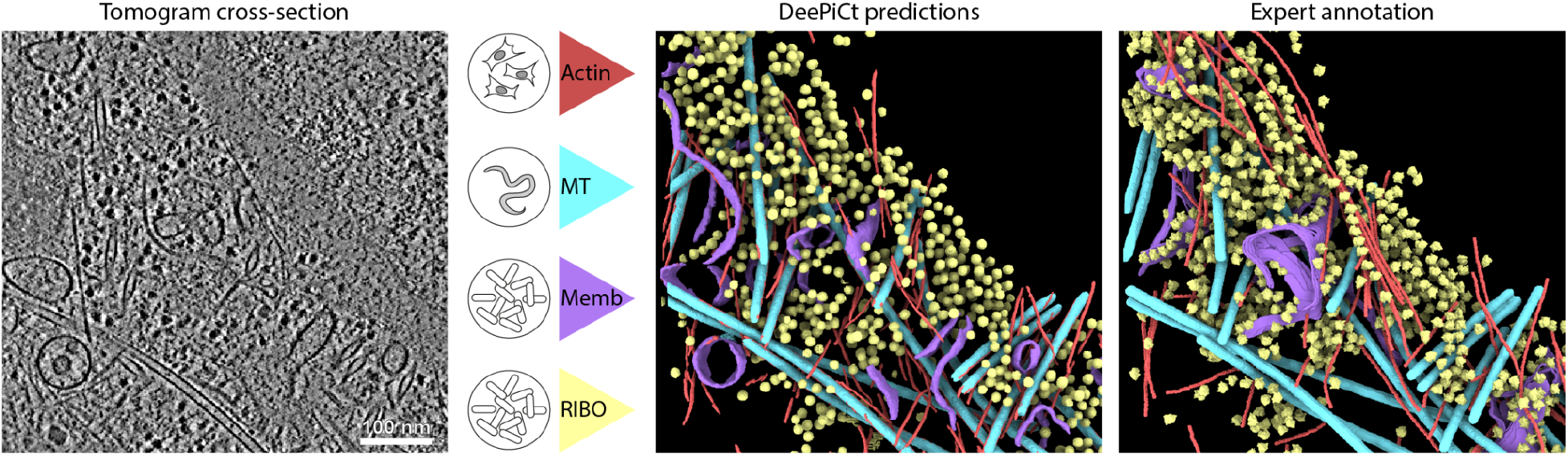
DeePiCt’s generalization across species. A dataset depicting a HeLa cell nuclear periphery1 is segmented by applying four independently trained DeePiCt networks. The results show the segmentation of actin filaments (red) trained on RPE-1 and MEF 3T3 tomograms, microtubules (MTs, cyan) trained on *C. elegans* tomograms, and cytosolic ribosomes (yellow) and membranes (purple) trained on *S. pombe* tomograms.

### DeePiCt predictions result in high quality subtomogram averages

In the case of FAS in the DEF dataset, fewer particles were predicted than in the ground truth (**Extended Data Tables 5, 6**). Nevertheless, the subtomogram average of the detected particles revealed the two half domes of the barrel-shaped type I FAS with applied D3 symmetry, consistent with the ground truth average and published structures of this fungal 2.6-megadalton heterododecameric (α6β6) complex from other yeast species (**Fig. 4c-d**, **Extended Data Fig. 8b-e**). The phosphopantetheine transferase (PPT) domains required for activation of FAS could be observed along the equatorial plane of the complex. In addition, three equatorial densities were resolved that could not be assigned based on published structures. Three densities inside each half dome connected to the central α wheel and in close proximity to the ketosynthase (KS) fit with the acyl carrier protein (ACP) of *Saccharomyces cerevisiae*^44^ (**Fig. 4d**, PDB: 2uv8) and *Pichia pastoris*^45^ (**Extended Data Fig. 8c**, EMD-12139). The ACP shuttles the growing acyl-chain between the different catalytic sites, and its localization has been suggested in connection with the activity of the whole multi-enzyme complex^44, 46–49^. Here, *S. pombe* FAS complexes within FIB-lamellae prepared from exponentially growing cells revealed a specific ACP site, which had not been observed by cryo-ET and thus in its native state before (**Fig. 4d, Extended Data Fig. 8, Supplementary Note 2**).

For ribosomes, the particle averages and numbers purely derived from DeePiCt predictions are comparable to the ground truth annotations in DEF (**Extended Data Tables 5, 6**; **Supplementary Note 2, Supplementary Fig. 4, 5**) complementing the performance analysis results described above. 3D refinements of these lamella-derived ribosomal particles from both ground truth and DeePiCt resulted in subtomogram averages with the nominal resolutions of 11 Å and 15 Å after multi-particle refinement in M^33^, respectively (**Fig. 4e**, **Extended Data Fig. 8f, Extended Data Tables 5, 6**). Hierarchical 3D classification revealed in both datasets a well-aligned class that was further refined in M to subnanometer resolutions of 9.3 Å and 9.4 Å for ground truth and DeePiCt predictions, respectively (**Fig. 4f-h**, **Extended Data Fig. 8i-k, Extended Data Tables 5, 6, Supplementary Note 2, Supplementary Fig. 4, 5**). This allowed the identification of tRNA occupying the P-site of the peptidyl transferase center (PTC) as well as the L1 stalk facing the E-site (**Fig. 4g-h**, **Extended Data Fig. 8j-k**).

### DeePiCt-predicted ribosomes reveal functional sub-populations

The large number of particles that DeePiCt localized in DEF in a high throughput manner allows for examining subpopulations of macromolecular complexes that are functionally distinct by downstream subtomogram analysis. Further classification of all DeePiCt-predicted ribosomes focused on the head of the 40S small subunit revealed a subset with additional densities close to the head and at the exit tunnel (**Fig. 4i**, **Supplementary Fig. 6**). This class was also detected in VPP and in DEF ground truth datasets (**Extended Data Tables 4-6**). In the DEF ground truth dataset, densities for P- and E-site tRNAs and the L1 stalk facing inwards are resolved (**Extended Data Fig. 9a-e**). The ribosome-bound ATPase eEF3 from *S. cerevisiae*^50^ fitted well into the additional head density (CC 0.8972, EMD-12062, **Fig. 5i**, **Extended Data Fig. 9 a-c**). During translation, this eukaryotic elongation factor facilitates binding of a new tRNA to the A-site of the ribosome via the ternary aminoacyl-tRNA–eEF1A–GTP complex^50, 51^. In addition, unassigned density at the ribosomal exit tunnel is observable for this class. Focused classification of all DeePiCt-predicted ribosomes at this location revealed a subtomogram average fitting the *S. cerevisiae* ribosome^52^ (CC 0.9657, EMD-1667) with the rRNA expansion segment ES27L in a particular configuration^53^ (CC 0.7938, PDB-3izd) connecting to an additional density close to the ribosomal exit tunnel (**Fig. 4j, Supplementary Fig. 7**). ES27L plays a role in translation fidelity and recruits enzymes to the ribosomal exit tunnel, such as the methionine aminopeptidase (MetAP) that co-translationally processes the nascent peptide chain^54, 55^. The nuclear export factor Arx1, which is released during cytosolic ribosomal 60S maturation in *S. cerevisiae*^56^, and its human homologue Ebp1, a translation regulator^57, 58^, similarly bind at locations of the observed extra density (**Fig. 4j, Extended Data Fig. 9h**). These binding factors recruit the flexible rRNA scaffold ES27L and cover the ribosomal exit tunnel with their MetAP-like folds. This structural class was also detected in DEF gt and VPP gt datasets (**Extended Data Fig. 9f-j, Extended Data Tables 4-5**).

Thus, DeePiCt predictions combined with structural analysis revealed functional subpopulations of ribosomes. Mapping them back into the tomograms revealed their 3D spatial distributions in relation to other macromolecules such as FAS and organelles in the cellular context (**Fig. 4b**).

### DeePiCt’s combination of organelle segmentation and high-throughput particle localization enables structural insights into ribosome-mitochondria association

One advantage of DeePiCt over state-of-the-art tools is that it allows studying particle populations within their cellular context, by recovering particles in close proximity to specific organelles selected from predictions by the 2D CNN. We recovered cytosolic ribosomes within 25 nm distance to predicted endoplasmic reticulum (ER) and mitochondria from 7 and 3 DEF tomograms, respectively. Using subtomogram analyses for both subsets, two classes could be separated, one with and one without a membrane density (**Supplementary Fig. 8-9**). In the first class, ribosomes associated with the membrane in a specific orientation, whereas they were randomly oriented in the second class. The latter class likely arises from the highly crowded nature of the *S. pombe* cytoplasm (**Fig. 4k-l**), which in the case of DeePiCt predictions in DEF data, is also affected by imperfect organelle segmentations (**Fig. 4b**). ER-bound ribosomes faced the membrane with the ribosomal exit tunnel in agreement with published structures from other species such as *S. cerevisiae*^1, 52, 59^ (**Extended Data Fig. 9 k-n, Extended Data Table 6**). Mitochondria-bound ribosomes, which posed a particular challenge to structural analysis in previous studies^59, 60^, were found to interact with the membrane at an angular offset of around 35° in comparison to the ER-bound ribosomes (**Fig. 4m-n**, **Extended Data Fig. 9o-r, Extended Data Table 6**). Interestingly, a density connecting the ribosome to the mitochondrial membrane was detected at the bottom of the large subunit in close proximity to the small subunit, but not to the ribosomal exit tunnel. It possibly represents the ribosome nascent chain complex (NAC) in contact with the mitochondrion receptor OM14^61–63^ which has yet to be structurally described.

In summary, ribosomes close to mitochondria and ER exhibit different interfaces with the respective membranes, facilitating specific protein nascent chain membrane insertion or transfer into the organelle^2, 52, 62^. These results highlight the power of DeePiCt for high-throughput particle localization and cryo-electron tomogram segmentation to rapidly gain new biological insights.

### Trained networks can be readily applied to other species

As a demonstration of the domain generalization potential of our workflow across species, we predicted segmentations of ribosomes, membranes and cytoskeleton structures (actin filaments and microtubules) in a published VPP HeLa cell dataset^1^ (**Fig. 5**). The application of the DeePiCt pipeline employed a manually generated mask using Amira^64^ of the cytoplasmic volume, excluding the nucleus. For the evaluation of our results, we computed the voxel-F1 score in comparison to the publicly available annotations (EMD-11992^1^). For ribosomes and membranes, we used the ribosome- and membrane-networks trained on the 10 VPP-tomograms from *S. pombe* described in previous sections. This resulted in a voxel-F1 of 0.55 for ribosomes and a voxel-F1 of 0.18 for membranes (**Extended Data Fig. 10a**). However, the available expert membrane segmentation for the dataset covered only ER membranes, which makes the voxel-F1 value less meaningful. Visual inspection of the predicted membrane segmentation revealed a good fit (**Fig. 5**).

Additionally we generated models for microtubules prediction, initially training a network for simultaneous actin and microtubule segmentations in 4 tomograms of cryo-FIB milled Human retinal pigment epithelial-1 (RPE-1) cells^65^. Due to the high preferential orientation of the cytoskeletal filaments in this data exhibiting stress fibers, the performance for microtubules’ segmentation was low resulting in a voxel-F1 score of 0.26 (**Extended Data Fig. 10a**). We therefore trained a second 3D CNN for microtubule segmentation on 11 tomograms from lamella of cryo-FIB-milled, dissociated *C. elegans* cells, containing 890 microtubules in a wide range of orientations. This new microtubule segmentation network achieved a voxel-F1 score of 0.83 (**Extended Data Fig. 10a**), exemplifying the importance of a training set with a high number of structures, as well as orientations to mitigate the effect of the missing wedge on the prediction performance.

Finally, we trained a dedicated network for predicting actin-filaments on 5 manually curated tomograms, 2 from RPE-1 cells and 3 from Mouse embryonic fibroblasts (MEF) 3T3 cells, that contained approximately 3,740 actin filaments (**Online Methods**). The actin filaments network shows a low voxel-F1 score of 0.10 in the HeLa cell tomogram (**Fig. 5**, **Extended Data Fig. 10a**), which is likely caused by the finer structure of actin in comparison to microtubules, the minimal set of orientations sampled in the small amount of training data, and to the fact that most training filaments are arranged in bundles, probably causing the CNN to learn this superstructure rather than individual filaments (**Extended Data Fig. 10b**).

Overall, these results show that the prediction of ribosomes and microtubules, representing large macromolecular complexes, is especially well-preserved across species and that the availability of diverse and well-annotated training datasets are crucial for good performance of DeePiCt. The results shown in **Fig. 5** highlight the use of trained networks from DeePiCt to segment novel datasets spanning different species.

## Discussion

Our DeePiCt workflow facilitates accurate and fast localization of diverse structures in cryo-ET of intact cells. The demonstrated high performance and the flexibility of the 3D CNN architecture offer the user a reliable tool for pattern recognition. This enabled us to detect particle species, such as FAS, that are low in abundance and have a less dense structural signature in comparison to ribosomes. The integration of structure segmentation (predicted by the 3D CNN) with the contextual information excludes false positives in the 3D localization of particles and structures segmentation task, and harnesses the cellular context to carry out spatial studies focused on regions of biological interest. This enabled us to investigate the 3D spatial distribution of macromolecules such as ribosomes in proximity to specific organelles (*e.g.* mitochondria) and to obtain structural insights with functional implications.

Here, we provide the first publicly available experimental cryo-ET dataset of 20 *S. pombe* tomograms under two microscopy acquisition settings (VPP and DEF), together with a high-quality comprehensive annotation of RIBO and FAS, membranes, organelle and cytosol segmentations. This constitutes the first gold-standard dataset in the field that is large enough for unbiased model training, which will enable benchmarking of current methods, and spur the development of future computational tools for unbiased data mining in cryo-ET data.

Subtomogram averaging of the annotated particles from either ground truth or DeePiCt predictions resulted in the first density maps of the *S. pombe* ribosome and fatty acid synthase. We observed differences in the analysis of subtomograms derived from either VPP and DEF tomograms, despite both being derived from wild-type *S. pombe* cryo-FIB lamellae (**Supplementary Note 2**). Due to the preservation of high frequency features in DEF data, a subset of well-aligned ribosomes could be identified and refined to subnanometer resolution densities from cryo-FIB lamellae. The obtained maps provide detailed structural insights; for example, the P-site tRNA occupying the PTC, which together with the elongation factor eEF3 detected close to the head of the 40S subunit in a subset of ribosomes, suggest active translation. Further classifications close to the ribosomal exit tunnel revealed ES27L in a specific configuration providing a platform for other factors to interact with the exit tunnel and a potentially protruding nascent chain. By utilizing the contextual information of the DeePiCt predictions, differences in relative membrane orientation of mitochondria- and ER-bound *S. pombe* ribosomes were detected. The observed connecting density between ribosomes and mitochondria likely highlights the not yet structurally characterized ribosome-bound NAC in contact with OM-14^61–63^ facilitating protein import into mitochondria^60, 61^.

The analysis of DeePiCt’s performance confirmed that data quality is an important factor in its predictive ability, as demonstrated in the case of NPC predictions in the DEF* dataset, which we classified into high and lower quality subsets based on lamella thickness and tilt-series alignment error. As the NPC has a high degree of structural flexibility on the subunit and diameter level inside cells^40^ it is a more challenging target than for example ribosomes and thus emphasizes the importance of higher quality data to achieve sufficient performance. The introduction of a pre-processing equalizing filter is also an important element of the pipeline that improves both the learning process during the training of 3D segmentation networks for particles with less dense print than the ribosome (*e.g.* FAS), and especially the generalization power across domains, including different microscopy acquisition conditions. Thus, amplitude spectrum equalization allows localizing macromolecules across different data acquisition types. In the case of the 2D network, although pre-processing did not improve model performance on the same-domain inference, it led to cross-domain improvements when training on VPP data and inferring on DEF data, or when training on both data types combined to segment organelles and cytosol. More elaborate tasks, such as prediction of individual organelle types, will likely require more training data or training of a dedicated 3D CNN with a tailored network architecture.

Our workflow allows easy adaptation to the segmentations of other structures, as demonstrated by the application of cytoskeleton segmentation networks. The networks show high-quality performance for microtubules in the HeLa cell dataset after training on data with broad orientation sampling of the filaments. Actin predictions revealed a low F1 score and therefore likely require more training data sampling different orientations of the structural features to improve performance. Altogether, the application of multiple segmentation networks to the HeLa cell dataset revealed that DeePiCt trained on datasets from different microscopes, species, and conditions lead to reasonably good results when the quality of data is high. Although more in-depth analysis is needed to study the specific limitations for the applicability of DeePiCt networks on other datasets, the results presented here constitute the first step towards conducting large-scale quantitative analyses for structural biology using cryo-ET on cells from different laboratories and open source datasets. In this sense, the generated ground truth annotations in this study are a starting point to provide the community with a resource to improve and further develop cryo-ET object segmentation and detection tools, which will ultimately enable further exploration of particles in their cellular context. Together with the trained networks and their demonstrated applicability to other species, as well as the flexibility of the DeePiCt workflow, the software harbors great potential for quantitative cryo-ET studies in the future.

## Acknowledgments

We thank Jenia Zagoriy and Wim Hagen for technical support, EMBL IT and especially Thomas Hoffman for computational support. I.D.T. and M.T.N were supported by a fellowship from the EMBL Interdisciplinary Postdoctoral Program (EI3POD) under Marie Skłodowska-Curie Actions COFUND. J.M. acknowledges funding from the EMBL and the European Research Council starting grant (3DCellPhase-760067).

## Author contributions

I.D.T., J.M. and J.Z. conceived the study. I.D.T. implemented the 3D CNN code, co-produced ground truth annotations, carried out numerical experiments to evaluate performance. S.K.G. prepared *S. pombe* cryo-FIB lamellae and acquired cryo-electron tomograms, co-produced ground truth annotations, carried out subtomogram averaging and structural analysis. A.M. implemented the 2D CNN and spectrum equalization filter, produced ground truth segmentations and conducted performance analysis of the 2D CNN. F.S. carried out training and prediction with 3D CNN networks, implemented code and created Colab Notebooks for prediction. C.Z. generated data and provided NPC annotations under the supervision of M.B. M.T.N. generated RPE1 and MEF 3T3 data and together with D.W.C.C. produced ground truth and trained networks for actin segmentation. F.T. generated data, ground truth and trained networks for microtubule segmentation. C.P. and A.K. provided valuable input for the design and generation of the CNNs. I.D.T., S.K.G., A.M., J.M. and J.Z. wrote the manuscript with input from all authors.

## Code and data availability

The DeePiCt code is assembled as two Snakemake pipelines (2D CNN and 3D CNN); the 3D CNN is implemented in the Pytorch framework while the 2D CNN is implemented in Keras, both in Python 3. The code, trained models, and link to the Google Colab notebook are available in the github repository https://github.com/ZauggGroup/DeePiCt.

Raw tilt series, tomograms, ground truth coordinates and segmentations are available via EMPIAR accession codes EMPIAR-10988 (*S. pombe*) and EMPIAR-10989 (RPE1). Subtomogram averages for *S. pombe* VPP and DEF ground truth annotations are available on EMDB (VPP gt: EMD-14404, EMD-14405, EMD-14406, EMD-14408, EMD-14409, EMD-14410, EMD-14411; DEF gt: EMD-14412, EMD-14413, EMD-14415, EMD-14417, EMD-14418, EMD-14419, EMD-14420; DEF DeePiCt predicted: EMD-14422, EMD-14423, EMD-14424, EMD-14425, EMD-14426).

Structural comparisons were performed with *S. cerevisiae* FAS (PDB: 2uv8^44^), *P. pastoris* FAS (EMD-12139^45^), eEF3 from *S. cerevisiae* (EMD-12062^50^), the *S. cerevisiae* ribosome (EMD-1667^52^) with the rRNA expansion segment ES27L in a particular configuration (PDB-3izd^53^), the nuclear export factor Arx1 bound to the 60S large ribosomal subunit *S. cerevisiae* (EMD-2169^56^), the human Ebp1 (EMD-1068^57^), *S. cerevisiae* ribosomes derived from extracted ER (EMD-3764^59^), and the ER-bound HeLa ribosomes (EMD-8056^1^). The large subunit (LSU, 60S) of a published *S. cerevisiae* 80S ribosome map (EMD-3228^66^) and the *S. cerevisiae* FAS map (EMD-1623^46^) were used as references for TM in *S. pombe* tomograms. The HeLa cell dataset is available via accession code EMD-11992^1^.

## Additional information

Supplementary Information is available for this paper.

## Online Methods

### Yeast cell culture

*S. pombe* K972 *Sp h-* wt haploid was kindly provided by C. Haering, originally from P. Nurse. Cells were recovered from frozen stock by streaking on YE5S agar plates (YES Broth, Formedium, 20 g agarose/L) and incubated at 30 °C for 1-3 days. Colonies were re-streaked on fresh YES agar plates and incubated 1-3 days at 30 °C. Single colonies were inoculated in 5 mL of YES medium (YES Broth, Formedium, PCM0302, FM0618/8573) and grown at 30 °C, 170 rpm overnight (NCU-Shaker mini, Benchmark). On the next day, cultures were grown to their log phase at OD600 of 0.5 - 0.6 and diluted beforehand in YES if necessary.

### Vitrification of yeast cells

A Leica EM GP (Leica Microsystems) was utilized to vitrify yeast cells at liquid nitrogen temperature. Yeast cells were either diluted to OD600 of 0.2-0.4 in YES medium or, following a prior wash step, in PBS containing 5 % or 10 % BSA as cryoprotectant. In the chamber, 4 µl were directly applied to TEM grids (Quantifoil R1/2, Cu 200 mesh, holey carbon or SiO2 film), which were glow discharged on both sides for 45 s (Pelco Easy glow) immediately before usage. Blotting from the back side of the support was performed for 1-2 s at 22 °C and 99% humidity. Cells were plunge-frozen in liquid ethane cooled by liquid nitrogen and transferred into grid boxes until further usage.

### Cryo-FIB of *S. pombe*

TEM grids containing vitrified yeast cells were clipped into an autogrid with a cut-out enabling cryo-focused ion beam (cryo-FIB) milling at shallow angles^67^. Mounted on a 45° pre-tilt shuttle, grids were transferred into an Aquilos Dual beam microscope (ThermoFisher Scientific). Prior to cryo-FIB milling at liquid nitrogen temperature, cells were sputter-coated with platinum for 10-15 s (1 kV, 10 mA, 10 Pa). Subsequently, a layer of organometallic platinum (reservoir at 28°C) was applied by opening the gas injection system (GIS) for 8 sec at a stage Z position of 3-4 mm below the coincidence point. In three independent sessions, three grids with 5 lamellae each were prepared at a milling angle of 15°. Agglomerations of several cells were micromachined in three steps of rough milling to a thickness of 5 µm at 1 nA ion beam current, 3 µm at 0.5 nA and 1 µm at 0.1 nA. Milling progress was visually monitored between each milling step with the scanning electron microscope beam (SEM, 10 kV, 50 pA). Fine milling was performed at 50 pA to a target thickness of 200 nm. To render the lamellae conductive for TEM imaging, they were sputtered with platinum for 5 s (1 kV, 10 mA, 10 Pa) and transferred into cryo boxes.

### Cryo-electron tomography of *S. pombe*

Autogrids containing lamellae were loaded into a Titan Krios (Thermo Fisher Scientific) such that the axis of the pre-tilt introduced by FIB milling was aligned perpendicular to the tilt axis of the microscope^68^. Cryo-electron tomography acquisition parameters are summarized in **Extended Data Tables 4-5**. Tomograms were acquired at 300 kV on a K2 Summit direct detection camera (Gatan) operating in dose fractionation mode and utilizing a Quantum post-column energy filter operated at zero-loss (Gatan). Magnification of 42,000 (EFTEM) with a calibrated pixel size of 3.37 Å was used for the ground truth VPP and DEF datasets and 3.45 Å for NPC data. Up to 14 tilt series were collected on a single lamella in low dose mode using SerialEM^69^. Starting from the lamella pre-tilt, images were acquired at 2.0 - 4.0 µm underfocus, in 2° increments within a range of + 58° to - 40° using a dose-symmetric tilt scheme^70^. The total dose was up to 120 e-/Å^2^ with a constant electron dose per tilt image. For the ground truth data, a set of 10 tilt-series were either collected with a 70 µm objective aperture or a Volta potential phase plate (VPP, Thermo Fisher Scientific^39^) with prior conditioning for 5 min. NPC data was collected as described in previous work^40, 71^, but at 1.5 - 4.5 µm underfocus, with 3° increments and an effective tilt range of + 50° to - 50°.

### Tomogram reconstruction

Tilt movie frames were aligned using a SerialEM plugin. Tilt series were filtered according to the accumulated electron dose by Fourier cropping using the mtffilter function in etomo (IMOD/BETA4.10.12^73^), and sorted by tilt angle using a python script. Four times binned tilt images (movie sums) were aligned in etomo (IMOD/4.9.4) software package^73^ using patch tracking (residual error 0.291 - 0.569 px) and tomograms were reconstructed via weighted back projection. Tomogram thicknesses measured in 3dmod ranged between 80 - 310 nm.

### Ground truth annotation for organelles, cytoplasm, and membranes in VPP and DEF

Cellular compartment segmentation was performed for both the VPP and DEF datasets. The annotations include 10 different organelle classes (mitochondria, vesicle, tube, ER, nuclear envelope, nucleus, vacuole, lipid droplet, Golgi apparatus, vesicular body, see **Extended Data Table 1**) and cytosol annotations, each identified through a unique numerical label to allow for selection of specific subsets of compartments.. Segmentations of the 10 VPP tomograms were performed manually in Amira^64^. These annotations were used to train a 2D CNN and to predict the initial segmentations in 10 DEF tomograms, which were first pre-processed with the spectrum equalization filter. These predictions were then manually corrected to obtain high-quality ground truth segmentations in DEF (**Extended Data Table 1**).

Membrane annotations were performed on the 10 VPP tomograms and 5 DEF tomograms. Initially, 5 VPP tomograms were annotated using Amira to manually segment every 2 to 3 slices, which were subsequently interpolated by means of Amira’s interpolation tool. These 5 VPP tomograms were then used to train a membrane segmentation 3D CNN, whose predictions on the remaining 5 VPP tomograms as well as in 5 DEF tomograms, were manually corrected in Amira.

### Ground truth particle annotation in VPP

RIBO and FAS were localized in 4-times binned tomograms (13.48 Å pixel size) in an iterative workflow. TM (RIBO) or manual annotations (FAS) (step 1, **Extended Data Tables 2-3**) were used for training 3D CNNs (step 2). CNN predictions were masked with a segmentation of the cytosol. Step 2 was repeated several times and each training round trained on the combined predictions of step 1 and the preceding round. Each round in step 2 consisted of three simultaneously trained networks sharing the same set of default hyperparameters (**Supplementary Tables 2-3**), except for the number of IF=4, 8, and 32, to provide a cumulative prediction less overfitted to the incomplete training data. The cumulative total predictions from steps 1 and 2 were then complemented by a round of visual inspection to eliminate false positives and manual picking to recover remaining false negatives (step 3, **Extended Data Tables 2-3**). Finally, the particle lists were cleaned for duplicates by applying elliptic distance constraints to the coordinates (**Supplementary Note 1**, **Extended Data Tables 2-3**).

In particular, for RIBO in the VPP dataset, the initial annotation (step 1) was performed as a combination of two sub-steps: step 1a employed TM performed with the pyTOM toolbox^36^. In detail, a 3D cross-correlation search over 1,944 Euler angle combinations was conducted using the large subunit (LSU, 60S) of a published *S. cerevisiae* 80S ribosome map (EMD-3228^66^) scaled to the corresponding pixel size, and a spherical mask with a diameter of 337 Å applied. The obtained coordinates corresponding to the top 2000-3000 highest cross correlation scores were manually revised in Gaussian filtered tomograms (sigma = 3) with tom_chooser (pyTOM toolbox^36^). This was followed by step 1b, which consisted of a non-exhaustive complementary manual annotation of particles missed by TM (**Extended Data Tables 2-3**). These initial ribosome coordinates served as input for step 2: three training rounds of CNNs were successively applied, as a global visual inspection confirmed that the false positive rate was low, while the fraction of true positives recovered was substantial. The resulting CNN predictions were manually revised in tom_chooser as described above. In step 3, remaining particles that were not detected by either TM or CNN were manually localized in either spectrum-matched or Gaussian filtered tomograms (sigma = 3) for up to 3 rounds in EMAN2 spt2_boxer^74^.

FAS in the VPP dataset was localized in step 1 manually in EMAN2, as TM with a published *S. cerevisiae* FAS^46^ map (EMD-1623) as reference failed (**Extended Data Tables 3**). In step 2 we performed only one round of training with these initial annotations and the resulting predictions were revised in EMAN2. Additional FAS particles were localized manually.

### Ground truth particle annotation in DEF

In the DEF dataset, the annotation procedure was the same as the one described for the VPP dataset, except that: 1) for RIBO the initial annotations relied on manually cleaned TM results and for FAS on incomplete manual picking, and 2) in step 2 of the RIBO ground truth construction, the first of the 3 rounds of CNN predictions used the DEF training data (from the TM results), while for the remaining 2 rounds the predictions were obtained using the models from the first and second rounds of RIBO VPP ground truth construction (trained only on VPP data, **Supplementary Table 2**). The visual inspection in step 2 was performed over the combined predictions of both the previously mentioned VPP networks and a round of DEF networks (IF=4, 8, 32) based on DEF TM results. After the final manual picking for missing particles (not present in either the initial TM results or the manually cleaned network predictions), we achieved comprehensive RIBO and FAS annotations (**Extended Data Tables 2-3**).

### Comparison of cryo-ET-derived particle numbers with proteomics

Copy numbers of ribosomes and FAS per cell were calculated with the ground truth annotations for an *S. pombe* cell with the assumption of 30 % cytosolic volume^75^ of a total of 150 µm^3^ average cell volume^76, 77^. It was also considered in the case of FAS that one fully assembled FAS complex is constituted by each six alpha and beta subunits.

### NPC manual localization

NPCs were manually localized as described previously^40^. In brief, coordinates and initial orientations of 354 NPCs were manually determined in 127 4-times binned, SIRT-like filtered^73^ defocus-only tomograms, as described earlier^71^. Two points located at opposing inner and outer nuclear membrane (INM-ONM) fusion points were manually picked for each NPC using IMOD. A third point was then chosen located at the INM-ONM fusion ring such that all three points define the horizontal plane of the NPC particle. The mean coordinates between the first two points were then used to assign the center of the NPC. The initial orientation of the NPCs was defined based on the normal vector to the defined NPC plane. A fourth point was chosen in the cytoplasmic side of the horizontal plane to assign the correct initial orientation and prevent false upside-down assignment of initial orientations. The annotated NPC data was divided based on quality criteria: 38 tomograms of quality 1 have a thickness below 300 nm, thus display optimal image contrast, and a tilt-series alignment residual error below 0.7 px. Quality 0 was assigned to 89 tomograms with 300 - 395 nm thickness and a residual error of 0.7 - 5.0 for tilt series alignments.

### Voxel-level representation of ground truth

As the final product of our ground truth generation process, we obtain voxel-based masks for membranes, cellular organelles, cytoplasm, and particles (RIBO, FAS, NPC). For RIBO and FAS, the lists of coordinates obtained were used to paste spherical masks (with radii of 10.78 nm for RIBO and 13.48 nm for FAS) centered on each of these points using a python script (Code and data availability). For the NPCs in DEF*, a subunit mask obtained by prior 3D averaging in novaSTA^78^, was pasted at each of the 8 subunit locations. The RIBO, FAS, membrane, organelle and cytosol masks in DEF and VPP along with the list of coordinates and respective tomograms are used in the subsequent training and performance analysis and are available on EMPIAR (Code and data availability).

### Cytoskeletal filaments segmentation

Microtubule and actin networks were trained on tomograms of Human retinal pigment epithelial-1 (RPE-1) cells previously described^65^. Segmentations of the individual cytoskeletal elements performed using the filament tracing function in Amira^79, 80^, followed by manual curation. Trained networks were applied to a tomogram depicting the nuclear periphery of a HeLa cell and compared against the corresponding segmentations (EMD-11992).

### Subtomogram analysis for ribosomes and FAS

Contrast transfer function (CTF) estimations, generation of 3D CTF models and subtomogram reconstructions were performed in Warp^32^. CTFs were first estimated in the sums of raw tilt movies and subsequently in the tilt series taking the tilt angles into account. Subtomograms containing ribosomes and their respective CTF models were reconstructed in volumes of 140³ pixels with a pixel size of 3.3702 Å and a particle diameter of 350 Å. Initial alignments of the subtomograms were performed in RELION version 3.0.7^34^ in 25 iterations of 3D classification, with the *S. cerevisiae* 80S ribosome (EMD-3228, low-pass filtered to 60 Å) as reference to generate an initial single class average. 3D refinements were performed with the resulting average as a reference. In the case of DEF data, this RELION-refined average was further refined in M^33^ to optimize particle poses, images and volumes warping to model non-linear deformations. Particles were re-extracted and hierarchical 3D classifications (25 iterations each) were performed in RELION. In the case of VPP data, hierarchical 3D classifications were performed directly after 3D refinements in RELION. Focused classifications were performed with binary masks indicated in the respective figures. Masks were generated from spheres produced in Matlab (MathWorks) in a cubic volume of 50 pixels per side and a radius of 20 pixels, scaled to a pixel size of 3.3702 Å and converted into binary masks resulting in a radius of around 130 Å using RELION, and placed relative to the area of interest using Chimera^81^.

FAS particles and CTF models were reconstructed in cubic volumes of 160 pixels per side with a pixel size of 3.3702 Å and a particle diameter of 400 Å. Initial alignments were performed in RELION in 25 iterations of 3D classification into a single class, and followingly refined with the 3D refinement option and applying D3 symmetry using the *S. cerevisiae* FAS map (EMD-1623^46^, low-pass filtered to 60 Å) as reference. Hierarchical 3D classifications (25 iterations each) were performed either after 3D refinements in RELION (VPP data) or after subtomogram re-extraction using the ribosome-optimized image and volume models in M (described above).

Final subtomogram averages of each particle class were obtained by 3D refinement and post-processing, including filtering to their respective resolutions determined by Fourier Shell Correlation (FSC) of two independently refined half maps at a cut-off of 0.143. Details of particle numbers and resolutions for each subtomogram average are summarized in **Extended Data Tables 4-6**. Visualization and calculation of cross-correlations (CC) between different maps and models, was performed with the UCSF ChimeraX package^82^.

### CNN Pre- and post-processing

#### Preprocessing

Tomograms are normalized to obtain uniform mean of 0 and variance of 1 in the frequency domain, prior to training. Additionally, the spectrum equalization filter is applied by matching the amplitude spectrum of the tomogram that is to be transformed to the target spectrum of one manually selected high-contrast VPP tomogram (Tomogram TS_001, **Extended Data** Fig. 1b). Extraction of spectra amplitudes is done using fast Fourier transform (FFT) followed by radially averaging amplitudes across the frequency domain. If the target tomogram’s Nyquist frequency is lower than the input tomogram’s, the target spectrum is padded with zeros to match the input spectrum’s size. Next, an equalization vector is created by dividing entry-wise the target spectrum by the respective input spectrum. The resulting equalization vector is converted into a rotational kernel and multiplied by the input tomograms in the frequency domain in combination with a sigmoidal-shaped low-pass filter to eliminate high frequency noise. After back transformation, the tomogram exhibits a similar contrast to the target tomogram (**Extended Data** Fig. 1). Tomogram intensities are additionally normalized to obtain uniform mean and variance. For the 2D CNN, tomograms and training segmentations are processed slice-wise into 2D tiles with a fixed size of 288x288 pixels (256x256 pixels and 16 pixels padding on each side). For the 3D CNN, tomograms are by default split into cubic patches of 64x64x64 voxels, and 12 voxels overlap in each dimension.

#### Post-processing (2D CNN)

The post-processing steps for the 2D network are focused on assembling the per-slice prediction into a 3D segmentation. To that aim, predicted tile segmentations are cropped on each side by 48 pixels to reduce artifacts around the edges, followed by reassembly into 3D stacks, with remaining overlapping areas being averaged. A 1D Gaussian filter is successively applied along the z axis in order to reduce single-slice false positives (**Extended Data** Fig. 2).

#### Post-processing (3D CNN)

After reassembly of the individual 64x64x64 voxel patches into the probability map resulting from the 3D CNN, the post processing step includes the clustering of the thresholded map (usually at threshold value of 0.5). According to user preferences, only clusters that lie within or close to a given mask consisting of an organelle or cytosol segmentation outputted by the 2D network, and whose size lies within a given range of numbers of voxels, are considered for the final prediction map (**Supplementary Note 1**). This ‘cleaning’ procedure allows adding *a priori* information on the size of the particles in question. Additionally, for macromolecular complex localization, the user has the option to export a list of coordinates of cluster-centroids, which represent the particles’ locations prediction.

#### Precision-Recall and Performance Analysis

##### Cross-validation approach

The analysis of performance for RIBO and FAS localization, as well as membrane segmentation, were done using a 3-fold CV scheme, where 8 out of the 10 VPP tomograms were used for training, and the remaining 2 VPP tomograms were used for testing. In the test for domain generalization, the same three networks (trained on the 8 VPP tomograms) were applied to the 10 DEF tomograms.

The analysis of performance in the NPC segmentation task was done in the dedicated DEF* of 127 tomograms. Importantly, in order to apply the DeePiCt pipeline, we employed a predicted segmentation of the nuclear envelope as the *region mask* in the *contact* mode during post-processing of the NPC prediction (**Supplementary Note 1**). The NPC prediction was studied under a 3-fold CV scheme, where each split consisted of roughly 42 tomograms. The nuclear envelope prediction was achieved through an independently trained 3D CNN on 18 tomograms belonging to the DEF* dataset for which a manual annotation of the nuclear envelope was available, and which were uniformly split across the 3 CV subsets.

For organelle and cytosol segmentation evaluation, model performance was evaluated on the voxel-level for the post-processed 2D CNN predictions produced by each CV fold’s individually trained model, using 5000 voxels picked randomly from each tomogram of the CV fold’s respective test set. Precision and recall (**Supplementary Note 1**) were computed at threshold values varying from 0 to 1 on the picked voxels to compute the area under the precision-recall curves (AURPC) for each tomogram.

#### Evaluation metrics

In the case of particles (RIBO and FAS), the localization evaluation was made directly from the predicted lists containing their centroids’ voxel coordinates. In the case of DeePict, the lists of particles’ coordinates are sorted according to a decreasing score (cluster size) value. For both RIBO and FAS, we defined a true positive when a predicted coordinate matched a ground-truth particle coordinate within a tolerance radius of 130 Å (**Supplementary Note 1**). The F1 score, which is calculated as the harmonic mean of the precision and the recall, provides a measure of the quality of the predictions. We chose this measure for easier comparison to previous studies for particle localization methods in the field of cryo-ET^26, 27, 29, 83^. Other metrics such as the area under the precision recall curve (AUPRC) generated by increasing length of particles’ list were disregarded as the ordering of the list (a function of cluster size) does not necessarily reflect the method’s confidence on positive classification.

The evaluation of the segmentation task is of a totally different nature, as the shape of the object should be taken into account voxel-wise. For the 2D CNN predictions of organelles and cytosol, we used the AUPRC score with varying threshold over the network’s output map (**Supplementary Note 1**). For Deepict’s predictions of NPC and membrane segmentation, we instead used the voxel-based F1 score (**Supplementary Note 1**), to maintain hegemony with DeePiCt particle prediction evaluations.

#### Hyperparameter tuning of DeePiCt and performance assessment

The previously described cross-validation schemes and evaluation metrics were not only used for the final assessment of methods’ performance (Fig. 3), but to study the effect of the different hyperparameters that could be set by the user in the case of the 3D CNN (IF, D, ED, DD, and BN). The experiments’ starting point was always the default hyperparameter combination *D* = 2, *IF* = 4, *BN* = 0, *ED* = 0, *DD* = 0 (**Supplementary Note 1**). In all cases, we tested first for different values of *D*, while keeping the rest fixed. Then, we fixed *D=D*,* where *D\** denotes the value for which best performance was achieved and repeated the same process for *IF*. We continued with each of the remaining parameters, *BN*, *ED* and *DD*. The results are reported in **Extended Data** Fig. 6. They allowed us to define the best hyperparameter combination for each of the structures studied, which are summarized in **Supplementary Table 1**.

In each case we combined the 3D CNN segmentation of the target structure with an appropriate *region mask* (**Supplementary Note 1**, **Supplementary Table 1)**: in the RIBO and FAS localization task, we used a cytosol prediction from the 2D CNN in *intersection* mode to eliminate false positives in organelles and the extracellular space; for membrane segmentation we used again the predicted cytosol from the 2D CNN, this time in *contact* mode to eliminate clusters that are in the extracellular space or within organelles; for NPCs we used a 3D CNN prediction of the nuclear envelope in *contact* mode to eliminate false NPC segmentations far from the nuclear boundary. For RIBO, FAS and membrane, both 3D CNN (for target structure prediction) and 2D CNN (for *region mask* prediction) in DeePiCt employed a 3-fold CV in the VPP dataset. In each fold, 8 tomograms were used for training and model selection while the remaining 2 tomograms were used for testing (**Extended Data** Fig. 6a**-c**). The NPC predictions, which we treat as segmentation tasks, were evaluated in the DEF* independent data selected for that purpose (**Supplementary Note 1** and **Extended Data** Fig. 6d**-f**). The *region mask* of predicted nuclear envelope segmentation was generated by a fixed 3D CNN trained on 18 of the 127 DEF* tomograms. These 18 tomograms were evenly distributed among the 3 splits of the DEF* dataset used for the corresponding 3-fold CV scheme employed for the hyperparameter tuning analysis of NPC segmentation by DeePiCt (**Extended Data** Fig. 6d**-f**). We used a 3-fold CV where the total 127 tomograms were split into 3 random subsets with roughly the same number of tomograms each. Each fold consists of two such sets as training data and the remaining one for testing. Throughout our experiments, high-quality DEF* tomograms (defined based on lamella thickness and tomographic alignment residual error as described above) consistently showed higher performance for the NPC segmentation task than the low-quality DEF* tomograms, regardless of architectural variations (**Extended Data** Fig. 6d**-f**). This hints at the importance of high data-quality in segmentation and localization tasks.

For the evaluation of the particle localization task, we compared the lists of coordinates of the particles in our ground truth data with the predicted coordinate lists. For both RIBO and FAS, we considered a true positive when coordinates coincide within a tolerance radius of 10 voxels (corresponding to approximately 130 Å). In this way, we computed the recall (the proportion of ground truth particles that were recovered by the method) and the precision (the proportion of predicted particles that were true positives) and reported the F1 score, which is the harmonic mean of precision and recall (**Supplementary Note 1, Supplementary Table 1**). For the structure segmentation task (such as the membrane or the NPC segmentation evaluation), we compared the ground truth masks with the DeePiCt predicted post-processed segmentation by calculating their voxel-wise precision and recall, and reporting the corresponding voxel-based F1 (voxel-F1) score, also known as Sørensen-Dice coefficient (**Supplementary Note 1**).

All experiments for the 3D CNN of the DeePiCt pipeline and DeepFinder were performed using NVIDIA 2080 Ti GPU, Cuda 10.0, Python 3 and Pytorch 1.3.1. For the 2D CNN, training was conducted using an NVIDIA 2080 Ti GPU and an NVIDIA V100S GPU used for performance evaluation, using CUDA 10.0, Python 3 and Keras 2.3.1 with a tensorflow 2.0.0 backend. Detailed lists of parameters used for the 2D and 3D CNN are available alongside DeePiCt’s source code (Code and data availability).

---

**Extended Data Fig. 1.**
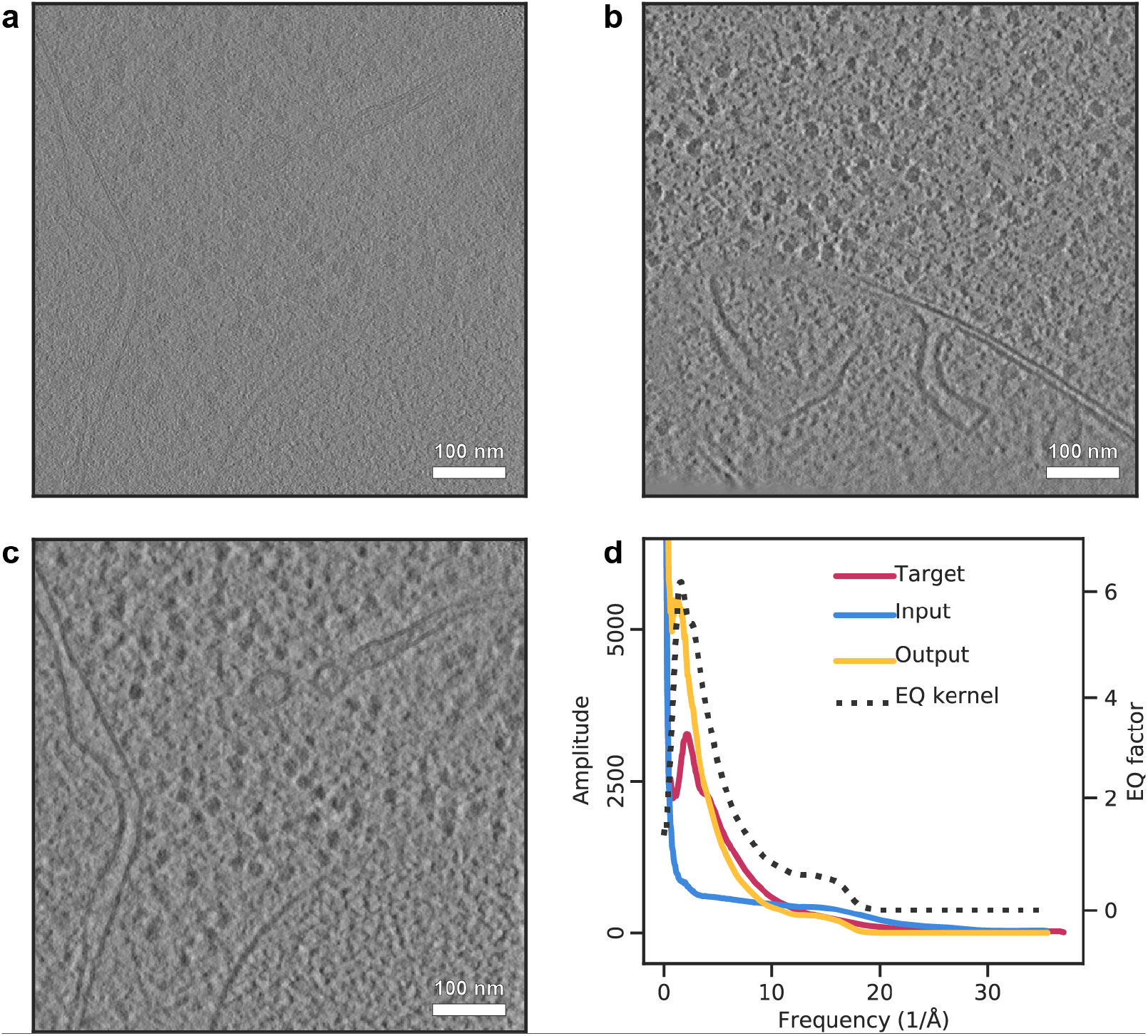
Pre-processing of input data by spectrum equalization. **a**. Input DEF tomogram with low signal-to-noise ratio. **b**. Target VPP tomogram with high signal-to-noise ratio. **c**. Output showing the tomogram in **a** after spectrum matching. **d**. Amplitude spectra of target, input and output tomograms, as well as per-frequency (1/Å) scaling factors of the rotational equalization kernel (y-axis cropped).

**Extended Data Fig. 2.**
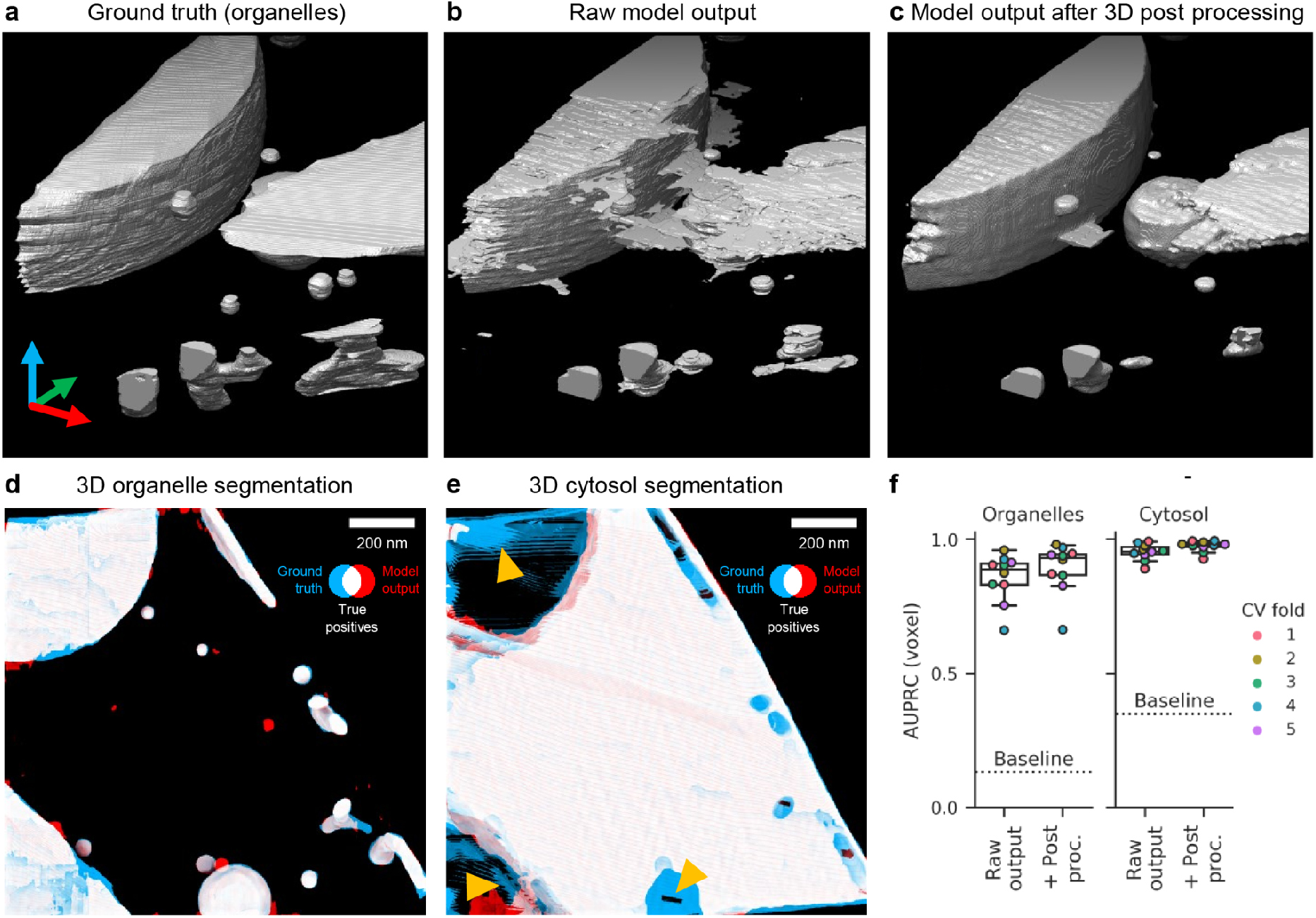
2D CNN organelle and cytosol segmentations. Isosurface view of organelle ground truth annotations (**a**), model output prior to (**b**) and after 3D post-processing (**c**). **d-f**. Organelle (**d**) and cytosol (**e**) predictions of the tomogram in **Fig. 1 b**. The 2D CNN’s segmentations (red) are overlaid with the manually created ground truth (blue), with overlapping regions (i.e. true positives) in white. Ground truth annotations in the top and bottom left corners in **e** (yellow arrowheads) are artifacts resulting from cross-slice interpolation in the manual annotation process. **f**. 3D post-processing improves performance of the 2D CNN as exemplified in (**a-c**).

**Extended Data Fig. 3.**
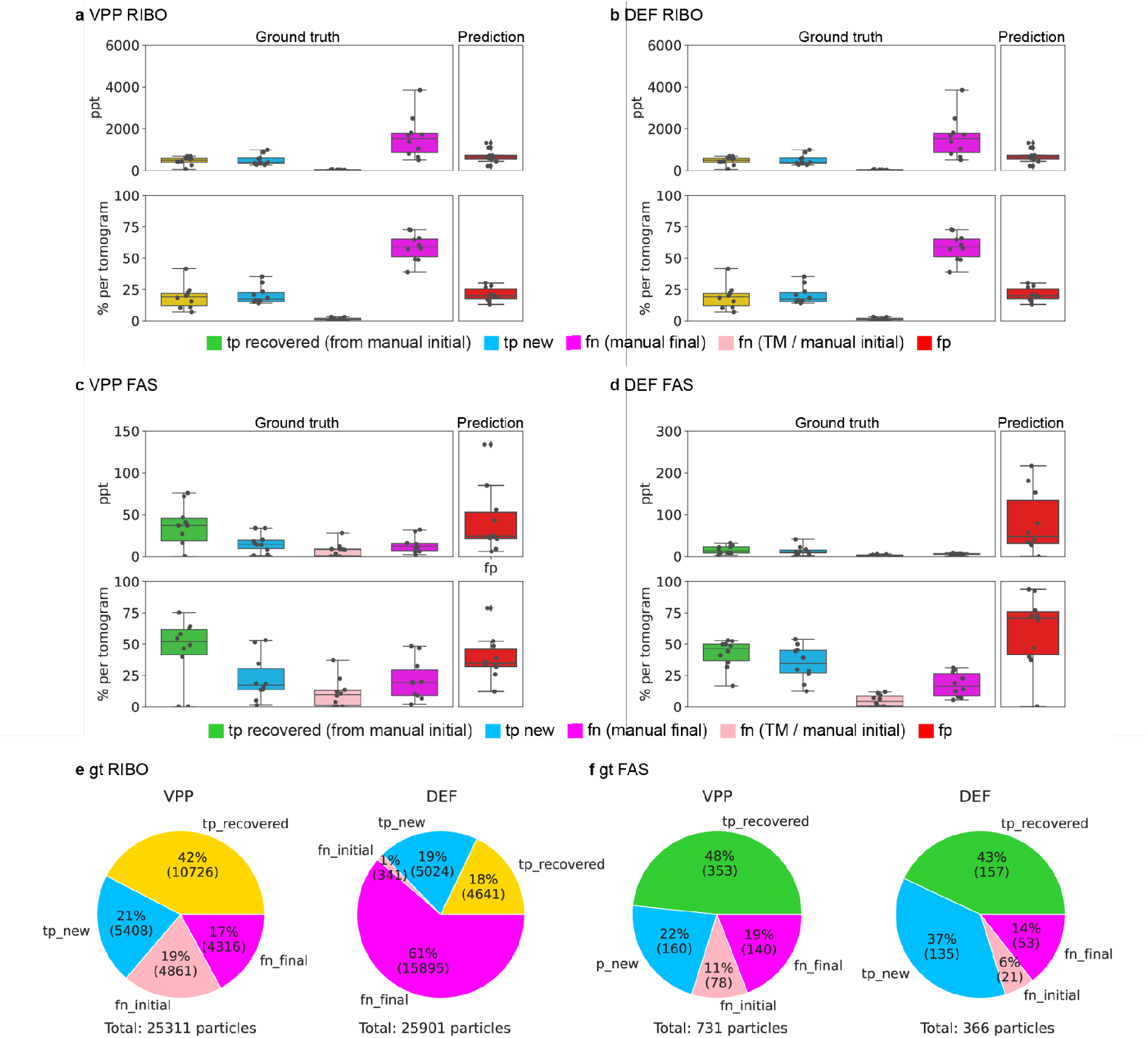
Details of the construction of ground truth for RIBO and FAS in VPP and DEF tomograms. **a**-**b**. Contributions of the steps applied during the construction of RIBO ground truth in the VPP (**a**) and DEF (**b**) datasets. Upper subplots show the absolute number of particles per tomogram (ppt): true positives from the initial annotations that were recovered in step 2 (yellow), newly identified true positives in step 2 (blue), particles that were in the initial annotation but that were not recovered in step 2 (false negatives from initial annotation; pink), unidentified particles from any of the two first steps, which were manually picked in step 3 (fuchsia), and total false negatives (red) in step 2. Lower subplots show the associated relative contribution, as a percentage of the ground truth per tomogram (ground truth panel) or as percentage of the predictions per tomogram in step 2 (prediction panel). **c**-**d**. Equivalent plots for FAS ground truth construction, where the only difference is that the recovered true positives originate from an initial manual annotation (green), as opposed to RIBO where we used TM additionally. **e-f**. Summary of the relative contributions in the RIBO (**e**) and FAS (**f**) ground truth across the 10 VPP and 10 DEF tomograms.

**Extended Data Fig. 4.**
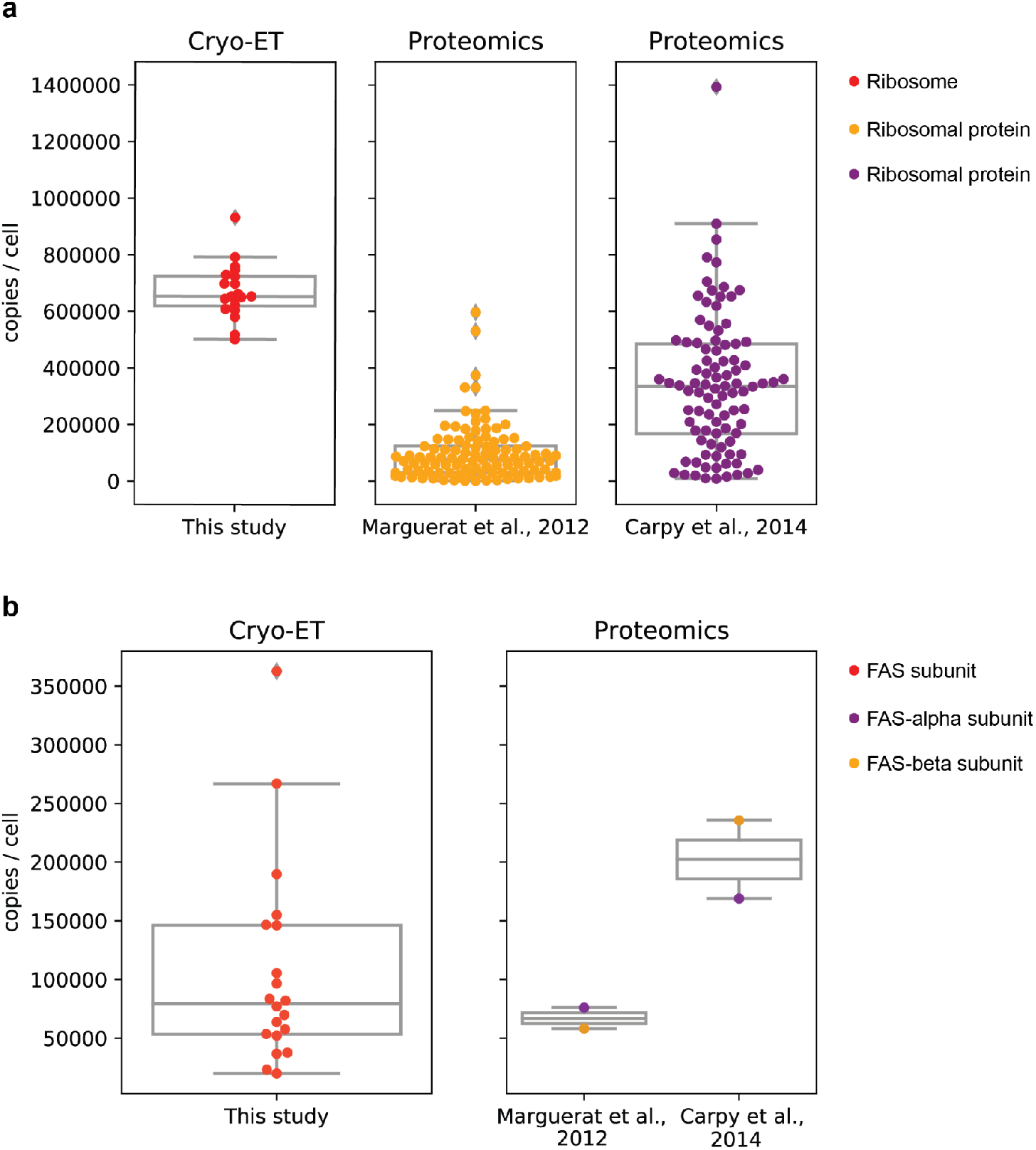
*S. pombe* ribosome and FAS complex abundances in cryo-electron tomograms in comparison to proteomics analysis. In this study (cryo-ET, left), fully assembled ribosomes (**a**) and FAS (**b**) were iteratively annotated in 20 ground truth cryo-electron tomograms and cytosolic concentrations calculated using the cytosol segmentations. Copy numbers per cell were calculated for an *S. pombe* cell with the assumption of 30 % cytosolic volume^1^ of a total of 150 µm^3^ average cell volume^2, 3^. **a**. With cryo-ET, an average of 671,303土 96,708 ribosomes/cell were annotated. Ribosome counts from individual tomograms are represented as individual points (red). Proteomics data of individual ribosomal proteins were derived from Marguerat et al.^4^ (yellow) and Carpy et al.^5^ (purple) and resulted in average of 97,795 土 99,298 and 343,511 土 244,226 ribosomes/cell, respectively. **b**. Each measurement displayed in the plot corresponds to 6 times fully assembled FAS counts per tomogram (the complex is constituted by six alpha and beta subunits). With cryo-ET, an average of 106,282 土 86,247 FAS subunits/cell were observed. Proteomics data of individual FAS proteins (subunit alpha in purple, beta in yellow) were derived from Marguerat et al.^4^ and Carpy et al.^5^ and resulted in average of 67,035土 12,600 and 202,263 土 47,086 FAS subunits/cell, respectively.

**Extended Data Fig. 5.**
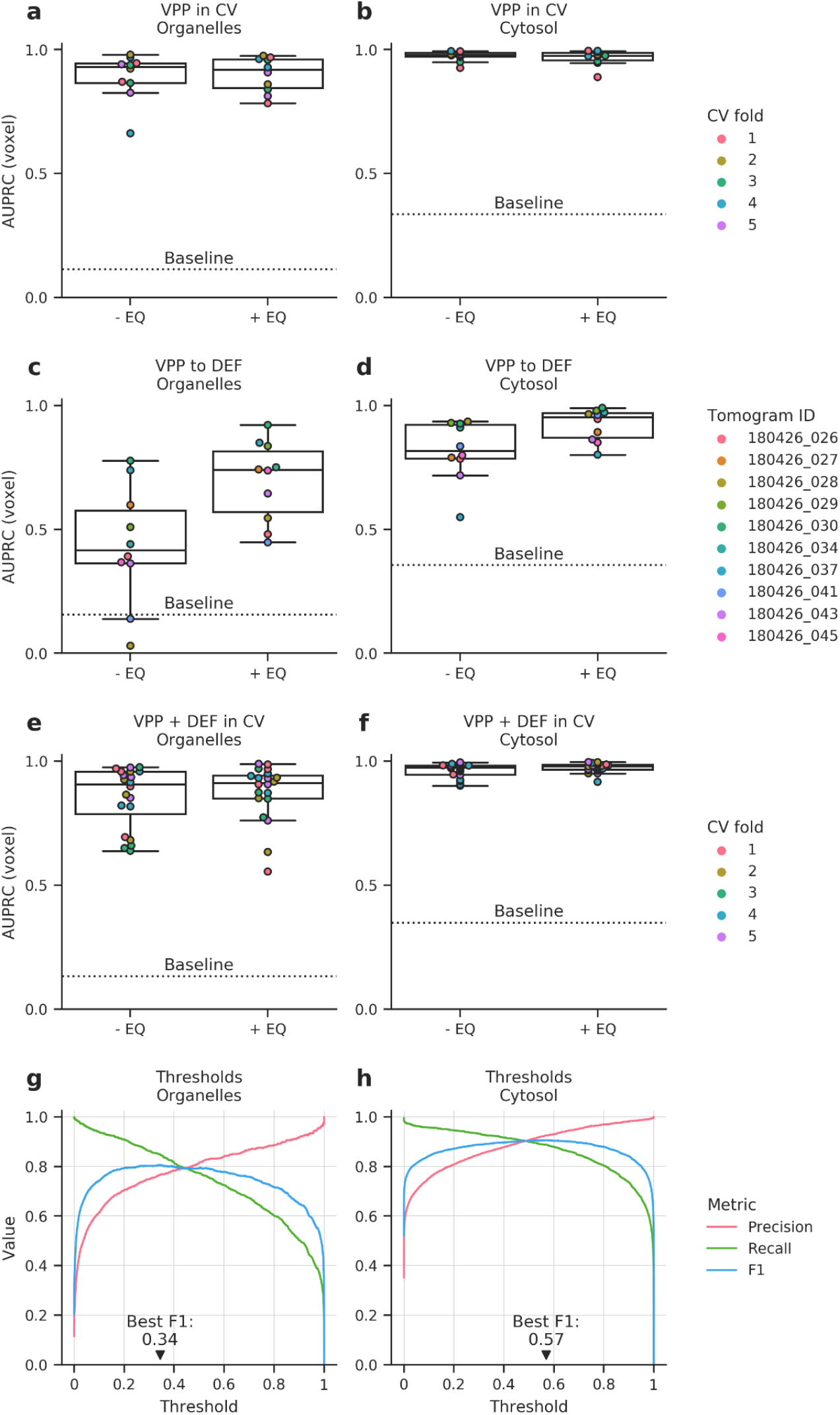
Effects of spectrum equalization on cross-validation and performance evaluation across domains of the 2D CNN. **a-f**. Performance was evaluated for 2D networks in three different scenarios (training on VPP in cross-validation, cross-domain generalization from VPP to DEF, training on VPP and DEF in cross validation), for either organelles or cytoplasm, without (-EQ) and with (+EQ) spectrum equalization. Areas under the precision-recall curves (AUPRCs) were computed by randomly picking 1000 voxels from each tomogram in test set (for the two cross-validation scenarios) or from the DEF tomograms (for the cross-domain tasks), while restricting evaluation to z-slices with any positive label. AUPRCs were computed after 3D post-processing. The baselines are defined as the fraction of positive labels within those z-slices and averaged across test tomograms. **g-h**. Precision, recall and F1-score of organelle (**g**) and cytosol (**h**) segmentation depending on probability threshold used. Scores were computed on the cross-validation results for the spectrum-equalized VPP tomograms.

**Extended Data Fig. 6.**
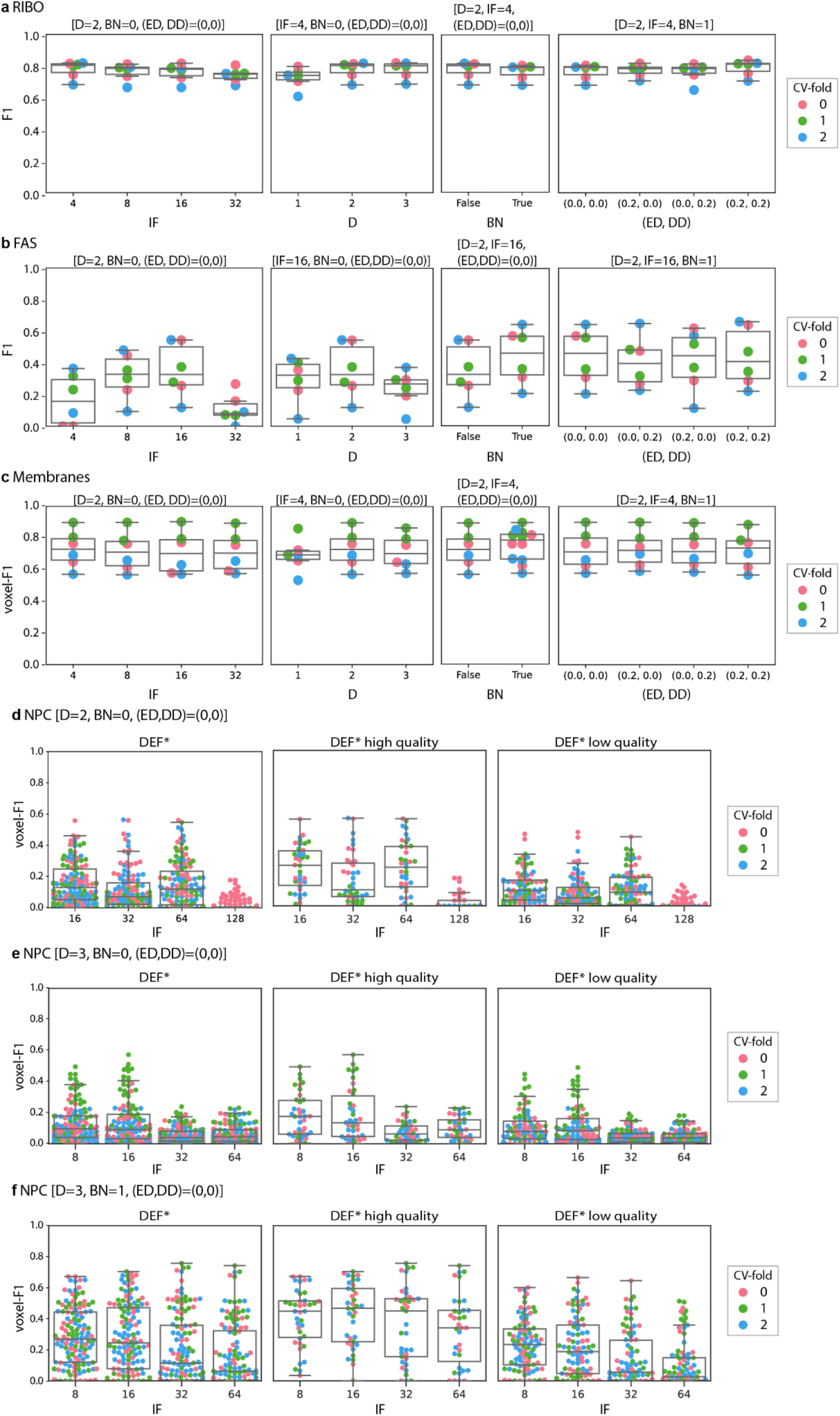
Hyperparameter tuning and performance analysis of 3D CNN for RIBO, FAS, membranes and NPCs. **a**-**c**. CV analysis for the performance of RIBO, FAS and membranes, respectively. In these experiments we follow an incremental search: from left to right, the starting subplot corresponds to the default configuration of the 3D CNN, D=2, BN=0, and (ED, DD)=(0,0), except for variable IF. The best value of IF (denoted by IF*) is fixed in the subsequent subplot to the right, where D is varied over 1, 2, and 3. The next subplot to the right compares the effect of not applying and applying the batch normalization layer (BN=0 and BN=1, respectively), under the best IF=IF* and D=D* from previous subplots. Finally, the right most subplot varies the encoder- and decoder-dropout parameters (ED, DD), with fixed best IF=IF*, D=D*, and BN=BN* from previous subplots. **d**-**f**. Results of hyperparameter exploration for NPC; in all DEF* tomograms (left), in the subset of high-quality DEF* tomograms (middle), and in the subset of lower quality DEF* tomograms (right). **d**. Shows the variations in IF, when D=2, BN=0, and (ED, DD) = (0,0); **e**. Variations in IF, when D=3, BN=0, and (ED, DD) = (0,0); **f**. Variations in IF, when D=3, BN=1, and (ED, DD) = (0,0). The summary of best hyperparameters’ combination is provided in **Supplementary Table 1**.

**Extended Data Fig. 7.**
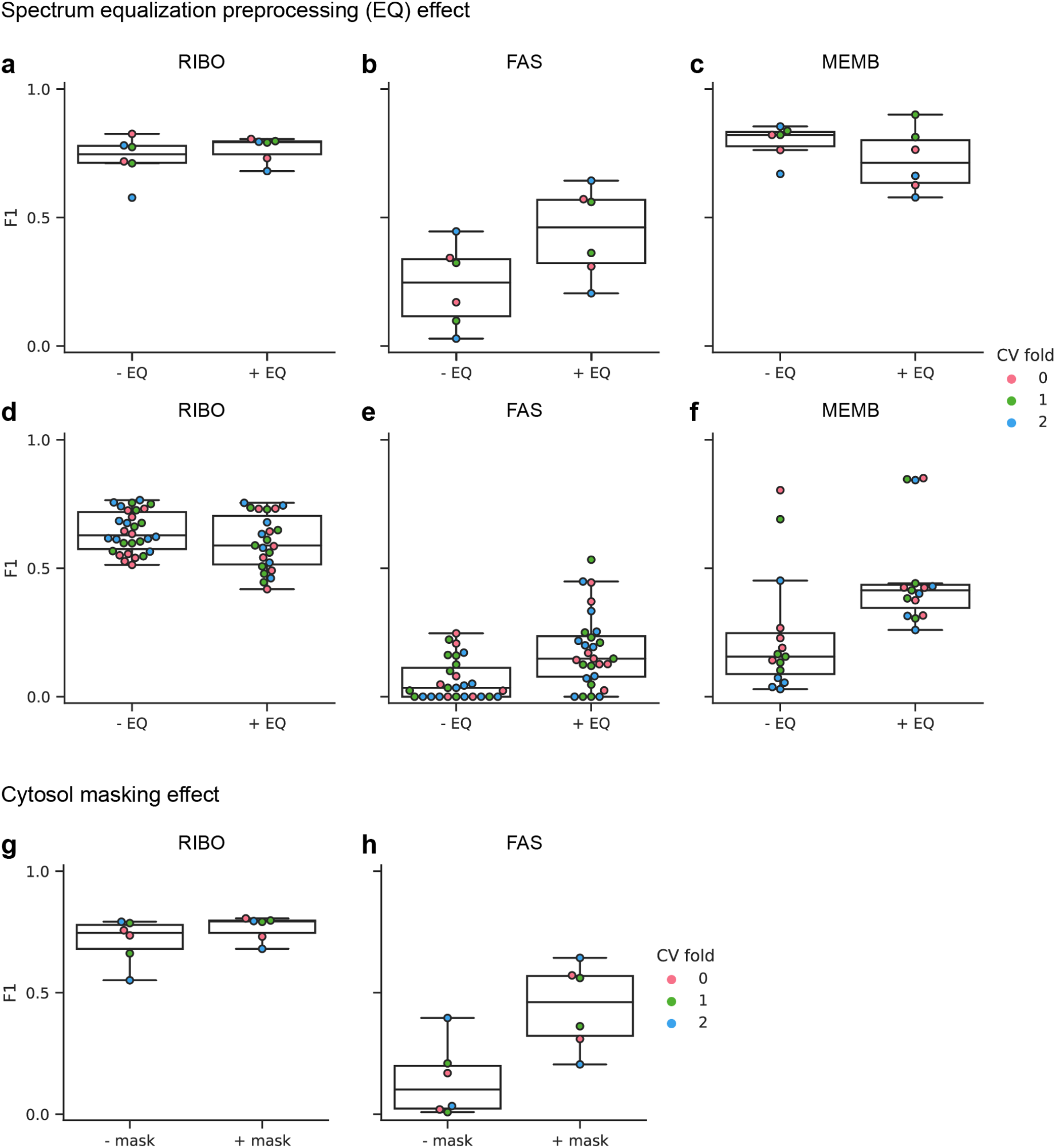
Effects of spectrum equalization and cytosol masking on DeePiCt performance. **a-c** Performance of the DeePiCt pipeline within the VPP domain for RIBO, FAS and membranes, without (-EQ) and with (+EQ) spectrum equalization pre-processing. The F1 score distribution shows that although for RIBO and membranes the spectrum equalization does not have a clear positive effect, for FAS it brings better results with a shift in median F1 score from 0.25 to 0.46. **d-f**. The same networks were used to test performance of the DeePiCt pipeline across the VPP and DEF domains (training on VPP and testing in DEF) for RIBO, FAS and membranes without (-EQ) and with (+EQ) spectrum equalization pre-processing. The F1 score distribution shows that while for ribosomes the spectrum equalization does not bring benefits, for FAS and membrane prediction across domains, its use brings a positive median F1 shift from 0.03 to 0.15 and from 0.16 to 0.41, respectively. **g-h**. Effects of employing a segmentation of cytosol as a *region mask* (+ mask) or not (-mask) during DeePiCt’s post-processing, for the localization of RIBO and FAS. The plots show the results under a 3-fold CV in VPP data, where the cytosol segmentation was obtained by a 2D CNN. **g**. RIBO localization does not show a significant improvement in F1 score when the cytosol masking is used. **h**. FAS results show a stronger difference in F1 score, with a shift of median F1 from 0.10 to 0.46.

**Extended Data Fig. 8.**
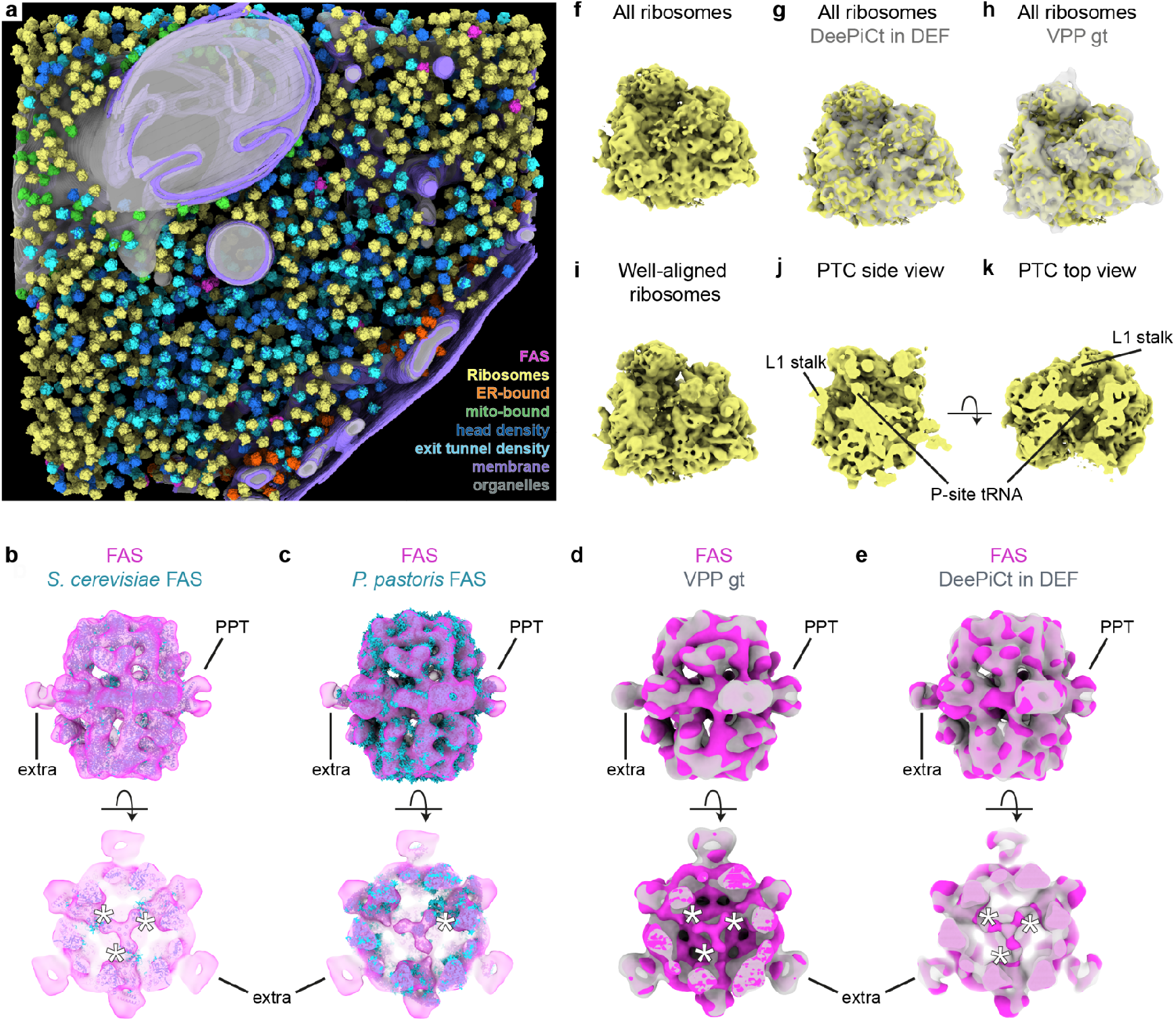
Ground truth annotations and 3D refinement of ribosomes and FAS in DEF tomograms. **a**. Complete annotation of ribosomes (yellow, orange, green, dark and light blue; sub-populations described in (**f**) and in Extended Data Figure 9), FAS (pink), membranes (purple) and organelles (gray) in the tomogram depicted in Fig. 4. Ribosome subclasses in the context of specific organelles were identified by a maximal distance of 25 nm to mitochondria (mito-bound, green) or ER (ER-bound, orange). Focused classification on densities close to the head of the 40S small ribosomal subunit or in proximity to the exit tunnel recovered ribosome subsets with head (dark blue) and exit tunnel density (cyan). **b-e**. FAS subtomogram averages (pink) fit of the *S. cerevisiae* X-ray structure (**b**, cyan) and the single particle cryo-EM density of *P. pastoris* (**c**, cyan) including the PPT domains. An extra density cannot be assigned. The cross section views close to the alpha-wheel reveal three densities fitting three and one ACPs of *S. cerevisiae* and *P. pastoris*, respectively (white asterisks, bottom). **d**. Overlay with FAS from VPP ground truth (gt) tomograms (gray) shows overall similarity, but the ACPs could not be resolved (white asterisks, bottom). **e**. FAS predicted by DeePiCt (gray) matches the DEF gt average with three ACP densities close to the alpha-wheel. **f-h**. Subtomogram average of all ribosomes (yellow) matches the DeePiCt-predicted (g, gray) and VPP-derived (h, gray) ribosome densities. **i-k**. Well-aligned ribosomes (yellow) detected by hierarchical 3D classification in RELION and refined in M (Supplementary Figure 3). **j-k**. Slicing through the average depicted in (**i**) at different axes reveals the PTC with a P-site tRNA and the L1 stalk facing the E-site.

**Extended Data Fig. 9.**
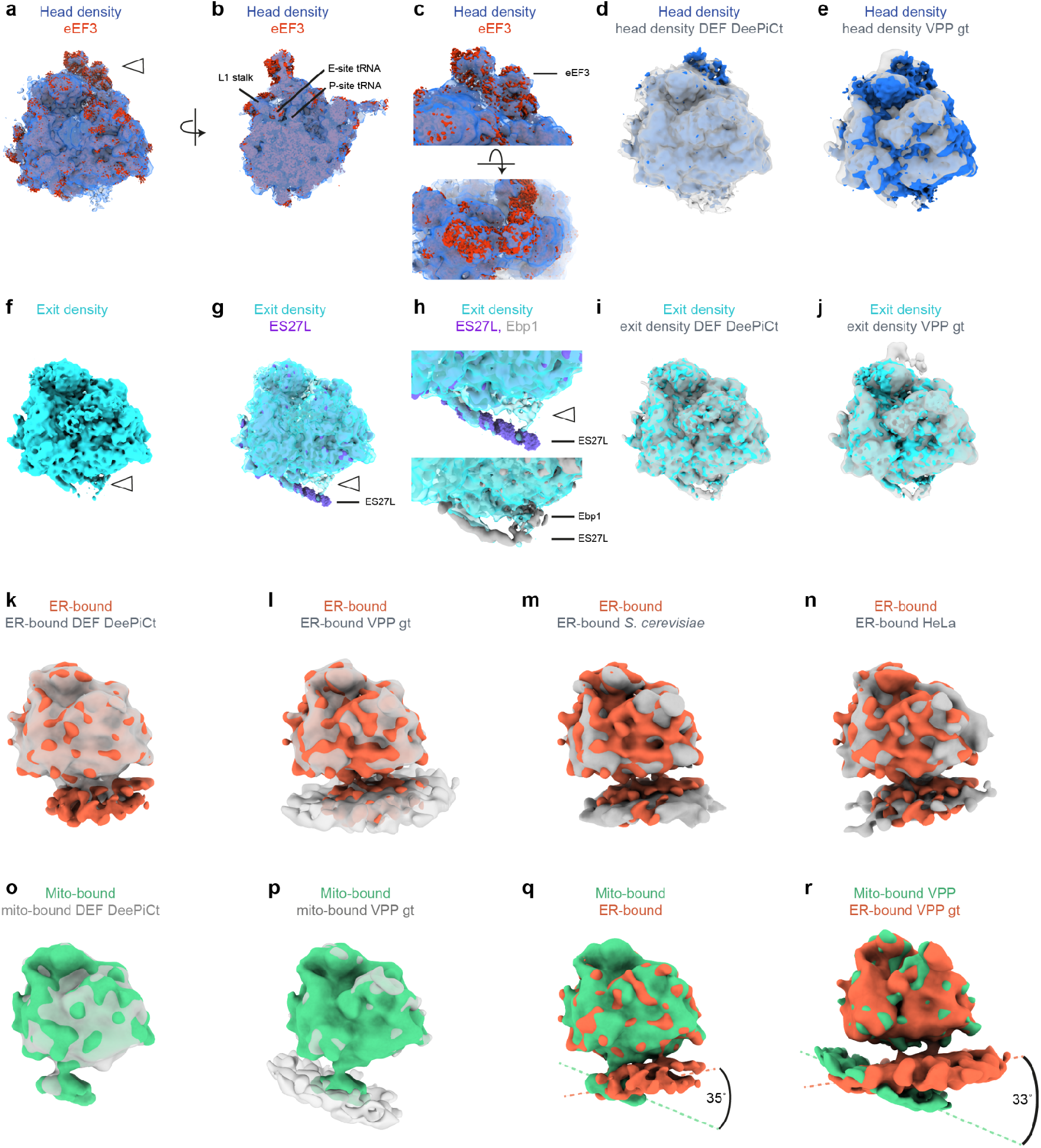
Comparison of ribosome subtomogram averages in DEF ground truth (gt). **a-e.** Subtomogram average of ribosomes (blue) with electron density in proximity to the head of the 40S small subunit (white arrowhead) which fits the ribosome-bound eEF3^6^ (CC 0.8972, red). **b**. Cross section of (**a**) showing the PTC with P- and E-site tRNAs and the L1 stalk facing inwards. **c**. Zoom into the additional head density fitting eEF3. **d-e**. Overlay with the same ribosome class from DeePiCt predictions in DEF (**d**, gray) and from VPP gt data (**e**, gray). **f**. Ribosome subset (cyan) with electron density in proximity to the exit tunnel (white arrowhead). **g.** Overlay with the S. cerevisiae ribosome^7^ (CC 0.9651, purple) and a specific conformation of the expansion segment ES27L^8^ (CC 0.7806, purple). **h**. Zoom into the exit tunnel area in (**g**) highlights the additional density connected to ES27L that colocalizes with for example the human Ebp1^9^ (gray, lower panel, CC 0.8495, EMD- 1068). **i-j**. Overlay with the same ribosome class from DeePiCt predictions in DEF (**i**, gray) and from VPP gt data (**j**, gray). **k-n**. Overlay of ribosomes within 25 nm distance to the ER (orange) with ER-bound ribosomes from DeePiCt predictions in DEF (**k**, gray), from VPP gt data (**l**, gray), with the published density of *S. cerevisiae* ribosomes derived from extracted ER^10^ (**k**, gray, CC 0.8972, EMD-3764), and the ER-bound HeLa ribosome^11^ (**n**, gray, CC 0.9105, EMD-8056). All densities show the same interface between ribosome and ER membrane. **o-p**. Overlay of ribosomes within 25 nm distance to mitochondria (green) with mito-bound ribosomes from DeePiCt predictions in DEF (**o**, gray), and VPP gt data (**p**, gray). All densities show the same interface between ribosome and mitochondria membrane. **q-r**. Comparisons between ER-bound (orange) and mitochondria-bound (mito- bound, green) subtomogram averages in DEF gt (**q**) and VPP gt (**r**) data reveal an angular offset of the membrane interfaces of 35° and 33°, respectively.

**Extended Data Fig. 10.**
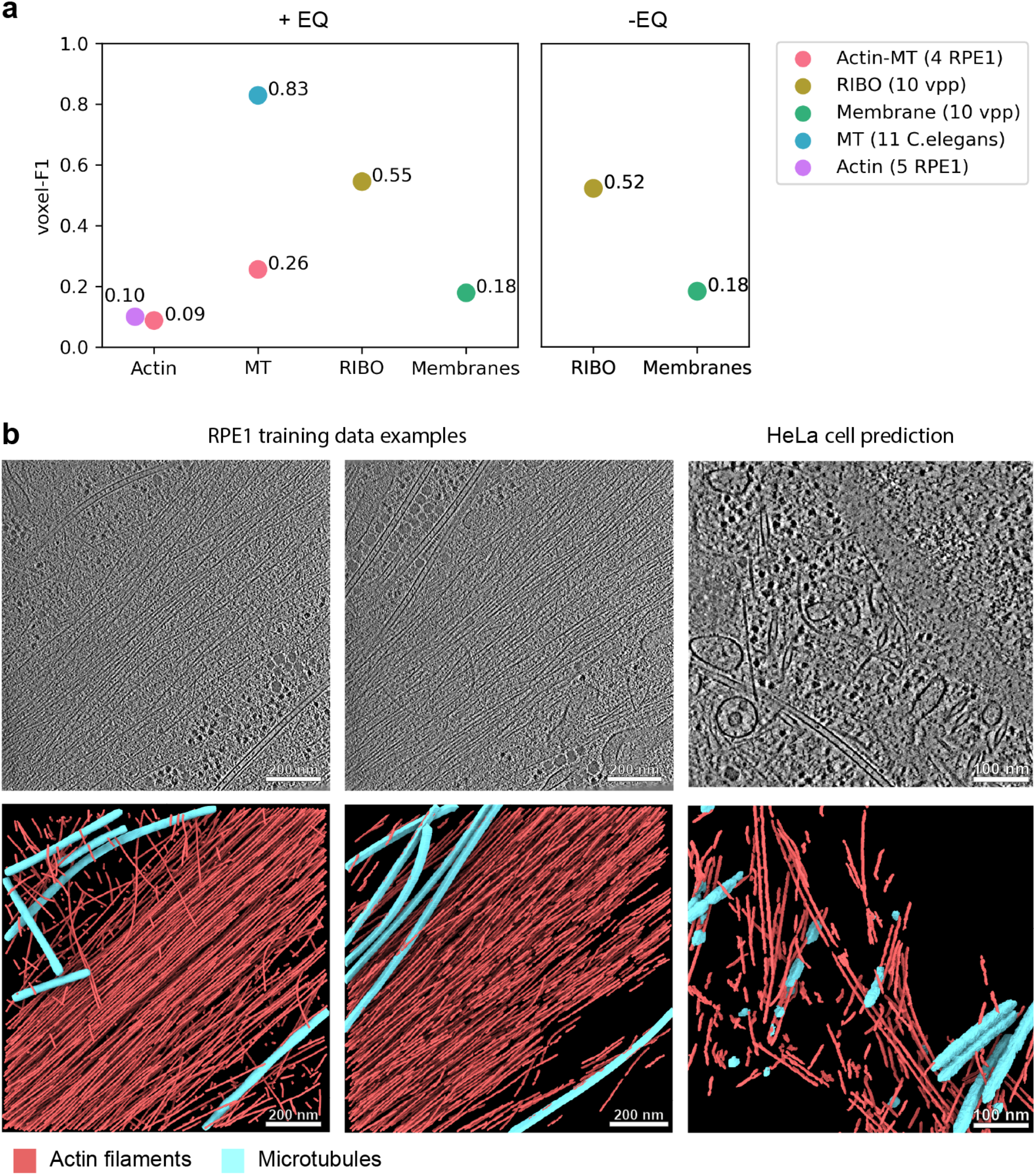
Domain generalization of different cellular structures and species. **a**. Performance results of five DeePiCt networks trained on a variety of cell types for the segmentation of different cellular structures used for predictions in a HeLa cell dataset^11^. Left: plot of performance scores when employing the spectrum equalization (+EQ) filter in both the training and on the HeLa cell data. The results correspond to segmentations of actin filaments and microtubules (MT) (trained on 4 RPE1 tomograms; pink), RIBO (trained on 10 VPP *S. pombe* tomograms; yellow-green), membranes (trained on 10 *S. pombe* VPP tomograms; aquamarine), MT (trained on 11 *C. elegans* tomograms; blue) and actin filaments (trained on 2 RPE1 and 3 MEF 3T3 tomograms with actin filaments at different orientations; light purple). Right: performance for RIBO and membrane without employing the equalization filter (-EQ) in the HeLa cell dataset. **b**. RPE1 training data examples (left and middle, tomograms on top, ground truth annotations at the bottom) and predictions (right) of the actin filament and MT models in the HeLa cell dataset. The training examples show 2 of the RPE1 tomograms^12^ used for this model. The performance of actin and microtubule predictions in the HeLa cell dataset is poor (voxel-F1=0.09 for actin and 0.26 for MT; **a** left) despite the high-quality training data (left). This is likely due to the high unidirectional orientation of the filaments in the training data (bundles in stress fibers).

## Supplementary Information Guide

### Supplementary Note 1

#### Architectural description of 2D and 3D CNN, post processing steps and evaluation metrics

The general architecture of the 3D and 2D CNN is based also on the U-Net architecture^1^ and consists of two consecutive symmetric paths: an encoder and a decoder path connected by skip-connections and a transitional base-layer (Fig. 1a), where:

a. The encoder path is a sequence of alternated convolutional blocks and max-pooling layers. In turn, the convolutional blocks are a series of convolutions (kernel size *k* = 3, stride *s* = 1 and zero padding *p* = 2) followed by a rectified linear unit (ReLU). In the case of the 3D CNN, optional batch normalization (BN) and dropout layers can be alternated (Fig. 1a). The max-pooling layer has fixed window size *w* = 2 and stride *s* = 2. In our implementations, the initial convolutional block has a variable number of initial filters (IF) in its first convolutional layer, which must be set by the user. In all cases, subsequent convolutional blocks duplicate the number of feature maps from the previous level. The number of downsampling layers plus 1, is referred to as the network’s depth (D), which is also a hyperparameter to be set by the user. It determines the size of the receptive field (*RF*) of the network (*RF* = 2 ∗ *D*). For the 2D CNN, the depth is a fixed value *D* = 5.
b. The base of the CNNs is the transitional convolutional layer between the encoder and decoder paths.
c. The decoder path is symmetric with respect to the encoder path, and is a sequence of alternated convolutional and upsampling (via transposed convolution) layers^2^, with kernel size *k* = 3, stride *s* = 1 and padding *p* = 1. For the 3D CNN, analogous optional layers can be alternated between convolutional and upsampling blocks (BN and dropout). The encoder and decoder paths’ associated parameters are denoted throughout the text as ED and DD, respectively. Essential to the U-Net architecture are the *skip-connections*, which consist of concatenating the feature maps from the encoder to the decoder path at every level. The last activation layer is in our case a sigmoid function, compatible with our choice of the Dice Loss function^3^. By definition, this is a multi-label approach, where single voxels may belong to multiple semantic classes, and which frees the user from defining additional weighting parameters to compensate for class imbalance necessary for other loss function choices such as categorical cross-entropy. In our experience, the use of Dice Loss provided better qualitative results in all cases. All models were trained using Adam Optimiser^4^. Application of these networks to unseen tomograms requires scaling of their pixel size to match (at least approximately) that of the training data.

### Data Augmentation

For the 2D CNN, tiles are randomly flipped and rotated in 90-degree increments during training to improve generalization.

Although it did not provide improved results in our hands (not shown), data augmentation for training the 3D CNN is available and optional. The number of times that each of the original training examples is augmented is given by a parameter *n_DA_* that can be set by the user. Each augmentation includes 4 different types of random volumetric transformations (applied to the original image and the labels, except in the case of noise), that we describe in what follows. Let A in *R*^2×2×2^ be a single channel cubic image of side-length *n* > 0. Then:

1. **Additive Gaussian noise.** Given a predefined modulation factor ε > 0, the Additive Gaussian noise transformation *G*_ε_ is defined by *G*_ε_(*A*) = *A* + ε*B*, where *B*_ijk_ ∼ *N*(0, 1) are independent and identically distributed random variables (i.i.d.r.v.), and *N*(*μ*, *σ*^2^) denotes a normally distributed Gaussian random variable with mean *μ* and variance *σ*^2^.
2. **Salt-and-pepper noise**. Given user-defined modulation factor *a* > 0 and a probability rate 1 > *p* > 0, the salt-and-pepper noise transformation is defined by *S_p,L_*(*A*) = *A* + *a* ∗ *B*, where *B*_ijk_ ∼ *U*(*p*, [−1, 1]) are i.i.d.r.v., and *U*(*p*, *I*) denotes a uniformly distributed random variable in a real interval *I*.
3. **Elastic transformation.** Given coordinate entries of A that are located at a *coarse* regular grid of thickness *n*_O_ > 1 are displaced by a random vector of i.i.d.r.v. uniform random variables. To achieve a smooth displacement vector field for all the points of the image *A* (i.e. on the *fine* grid), we generate the elastically transformed image as the interpolated vector field (via polynomial interpolation of order *k* > 0, defined by the user).
4. **Random rotation**. Given a probability of rotation *p* ∈ (0, 1) and angle range *α* ∈ (0, 180), the image A is rotated with a probability *p* with and angle in [−*α*, *α*], with respect to the *z* −axis.

#### 3D CNN default hyperparameters

A default, starting point set of hyperparameters values for the 3D CNN for this study: *D* = 2, *IF* = 4, *BN* = 0, *ED* = 0, *DD* = 0, and *n*_01_ = 0. We used them to later, through testing variations on them, study the optimal combinations of hyperparameters in the different segmentation and localization tasks (**Supplementary Table 1**).

#### 2D CNN post-processing

Post-processing of the 2D CNN includes the reassembly of predicted tile segmentations, by cropping 48 px on each side to reduce artifacts around the edges. Areas of sets of tiles still overlapping are averaged after cropping. In order to remove inter-slide discontinuities, a Gaussian filter with parameter σ > 0 (default σ = 5) is then applied along the z axis.

#### 3D CNN post-processing

1. Thresholding of the final probability map outputted by the last layer of the neural network (default threshold is the unbiased value 0.5, but is set by the user in the pipeline’s configuration file).
2. Clustering of the resulting thresholded map via the *connected component labeling label* algorithm, using the function *morphology.label* of the *scikit-image* python library^5^.
3. Integration with other segmentation maps, *e.g.* with the output of the 2D CNN. Importantly, a subsequent selection of clusters is made with respect to a *region_mask* provided by the user, which consists of an auxiliary binary image defining a tomographic region of interest (Fig 1b). We then allow 3 types of options (*intersection*, *contact*, or *colocalization*) for employing the *region_mask* through the corresponding *contact_mode* parameter, as follows: a. *contact_mode: intersection* The region_mask is used to directly mask the output of point 2, by considering only the overlapping region, *e.g.* including particles that are within an organelle’s mask.
b. *contact_mode: contact* Only clusters that have voxels in common with the region of interest are kept, *e.g*. selecting NPC on the nuclear envelope.
c. *contact_mode: colocalization* Only clusters whose centroid is located at a given distance (*colocalization_radius*) from the region of interest are kept, *e.g.* particles within a distance to an organelle. At this step, only clusters within the size range [*m*, *M*] are kept, where 0 ≤ *m* < *M* ≤ ∞ are defined by the user in the configuration file.
4. If the *calculate_motl* parameter is set to *True*, a list of the clusters’ centroids resulting from previous steps is saved as a 4-columns comma separated values (csv) file, where the first column is the score (cluster size), and the three remaining columns indicate the (*x*, *y*, *z*) centroid voxel coordinates. The list is sorted according to decreasing-score.

#### Segmentation evaluation

Given a 3D cellular segmentation prediction *I*_*pred*_ (resulting either from the 2D or the 3D CNN), its quality can be voxel-wise evaluated, by comparison to a ground truth binary image *I_GT_*, according to different metrics:

1. Voxel-based precision (P) is defined by:

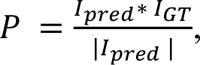

where the bars |indicate the number of entries different from 0. Thus, P is the proportion of voxels in the predicted segmentation that *truly* belong to the wanted structure.
2. Voxel-based recall (R) is defined by:

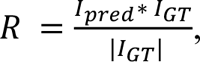

thus indicating the proportion of voxels of the ground truth segmentation that were correctly recovered by the prediction.
3. Area under the precision-recall curve (AUPRC). Since *I*_*pred*_ is dependent on the threshold value 0 ≤ *thr* ≤ 1 applied to the probability map, both precision and recall depend in ii as well, *P* = *P*_*thr*_ and *R* = *R*_*thr*_. The area under the precision-recall curve (AUPRC) is the area under the parametrized curve (*P*_*thr*_, *R*_*thr*_) is a widely used tool to summarize the power of the method.
4. Voxel-based F1-score (also known as Sørensen-Dice coefficient). A common metric to calculate the performance of the method by combining the P and R values is simply through the F1 score, defined by their harmonic mean:

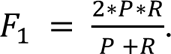 Notice that here no account of the dependence with respect to the *thr* value is used.

#### Particle detection evaluation

In the case of particles as discrete objects that are small enough to be defined by their coordinates, the evaluation of the object detection task requires the following definitions:

1. Given a sorted list of predicted particles’ centroids, one by one (in that order) they are either classified as false positives (fp) or true positives (tp).
2. A predicted particle coordinate is considered a true positive if and only if given a pre-defined tolerance radius r (*r* = 10 vox for both RIBO and FAS), there exists a ground-truth coordinate not previously matched to any other predicted particle.
3. Then the coordinate-based precision (p), recall (r), F1 score and area under the precision-recall curve (AUPRC) are defined by: 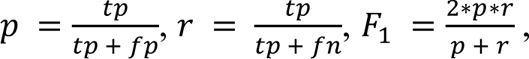, and AUPRC is the area under the curve defined by the (*p, r*) points when parametrized with respect to the cluster number (from 0 to all).

#### Merging several lists for ground truth construction

To account for possible duplicates when merging lists of particle coordinates from different methods (e.g. manually-, TM- and CNN-originated lists), we integrate them by restricting the distance between centroid coordinates, *i.e.* with the constraint *d*(*p*, *q*) > 1, where the elliptic distance *d*(*p*, *q*) is defined by:

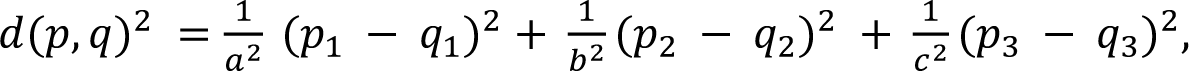

for *a* = *b* = 9, *c* = 15, and where *p* = (*p*_p_, *p*_E_, *p*_z_) and *p* = (*p*_p_, *p*_E_, *p*_z_).

The choice of the elliptic distance coefficients takes on account both the effect of elongation along the z-axis in the cryo-ET images and the size of the particle in voxels of the tomogram (1 vox = 13.48 Å).

#### Supplementary Tables

**Supplementary Table 1:** Selected hyperparameters of DeePiCt for varying cellular structures. Performances were calculated by cross-validation (CV) within the same domain, or in the case of microtubules and actin networks, in evaluations for generalization across cellular species with respect to expert annotations in the HeLa cell tomogram ("). DeePiCt networks were applied to the test datasets with the following *region mask* and *contact mode* combinations: cytosol and intersection for ribosome and FAS; cytosol and contact for membranes; nuclear envelope and contact for NPCs; cytosolic volume and intersection for microtubules and actin filaments.

| Cellular feature | Method | Depth (D) | Initial filters (IF) | BN | Median Performance | Training dataset | Testing dataset |
| --- | --- | --- | --- | --- | --- | --- | --- |
| Cytoplasm | 2D CNN | 5 | 16 | 1 | AUPRC=0.97 | VPP | VPP |
| Organelles | 2D CNN | 5 | 16 | 1 | AUPRC=0.92 | VPP | VPP |
| Ribosomes | DeePiCt | 2 | 4 | 1 | F1=0.79 | VPP | VPP |
| FAS | DeePiCt | 2 | 16 | 1 | F1=0.46 | VPP | VPP |
| Membranes | DeePiCt | 2 | 4 | 1 | voxel-F1=0.79 | VPP | VPP |
| NPCs | DeePiCt | 3 | 16 | 1 | voxel-F1=0.24 | DEF* | DEF* |
|  |  |  |  |  | voxel-F1=0.47 | DEF* | DEF* (high quality) |
|  |  |  |  |  | voxel-F1=0.19 | DEF* | DEF* (lower quality) |
| Microtubules | DeePiCt | 2 | 4 | 1 | voxel-F1=0.83<sup>†</sup> | VPP | HeLa cell |
| Actin | DeePiCt | 3 | 8 | 0 | voxel-F1=0.10<sup>†</sup> | VPP | HeLa cell |

**Supplementary Table 2:** Specifications of three 3D-CNN training and prediction rounds performed for ribosome ground truth construction. Each of the rounds in step 2 of the RIBO ground truth construction consisted of training 3 different 3D CNNs with default parameters except the IF number (**Supplementary Note 1**). The first training was performed on manually curated TM results of the corresponding dataset (VPP or DEF). For VPP, in the second round, the 3D CNN were trained on aggregated TM results and first round predictions; whereas the third CNN round was trained on all previous aggregated results. For DEF, the second and third round of 3D CNNs corresponded to the VPP 3D CNNs of first and second rounds, respectively.

| training round | Training dataset | Prediction dataset | Training set annotations | Initial filters | output |
| --- | --- | --- | --- | --- | --- |
| 1 | VPP | VPP | TM annotations | IF=4, 8, and 32 | vpp_output_round1 |
| 2 | VPP | VPP | TM annotations + vpp_output_round1 | IF=4, 8, and 32 | vpp_output_round2 |
| 3 | VPP | VPP | TM annotations + vpp_output_round1 + vpp_output_round2 | IF=4, 8, and 32 | vpp_output_round3 |
| 1 | DEF | DEF | TM annotations | IF=4, 8, and 32 | def_output_round1 |
| 2 | DEF | DEF | TM annotations + def_output_round1 | IF=4, 8, and 32 | def_output_round2 |
| 3 | VPP | DEF | TM annotations + vpp_output_round1 + vpp_output_round2 | IF=4, 8, and 32 | def_output_round3 |

**Supplementary Table 3:**
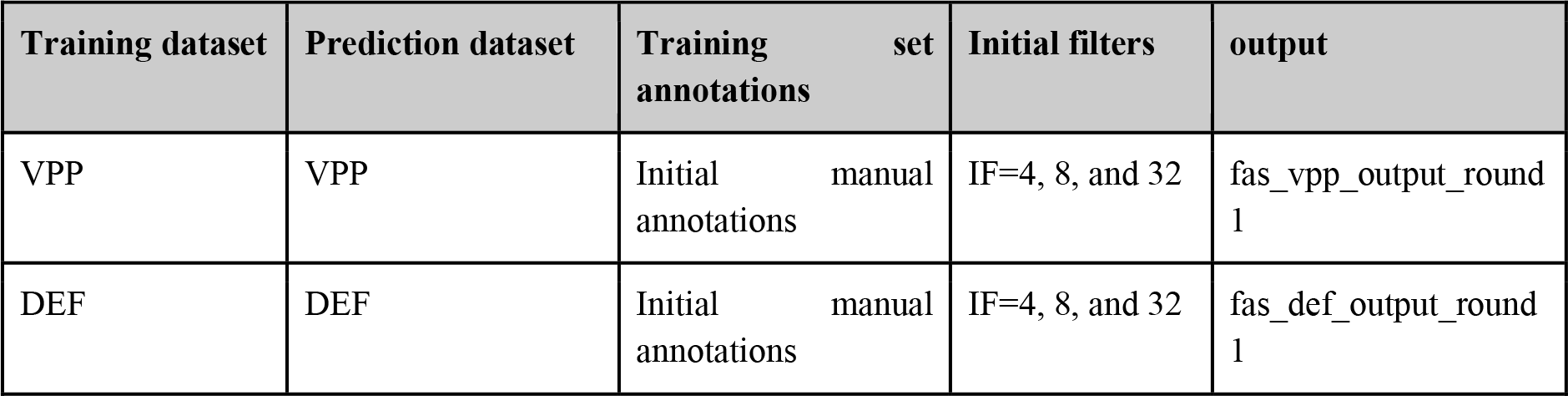
Specifications of 3D-CNN training and prediction rounds performed for FAS ground truth construction. Step 2 in FAS ground truth construction consisted of a single round of 3D CNN training, performed separately per dataset type (VPP and DEF). The round consisted of 3 networks with default parameters (**Supplementary Note 1**) except for the IF number, set to IF=4, 8, and 32. The networks were trained on an initial incomplete manual picking.

| Training dataset | Prediction dataset | Training set annotations | Initial filters | output |
| --- | --- | --- | --- | --- |
| VPP | VPP | Initial manual annotations | IF=4, 8, and 32 | fas_vpp_output_round1 |
| DEF | DEF | Initial manual annotations | IF=4, 8, and 32 | fas_def_output_round1 |

### Supplementary Note 2 Considerations for structural analysis following CNN predictions under different imaging conditions

Cryo-ET datasets of *S. pombe* utilized in this study were derived from cryo-FIB lamellae of wild-type, native, yeast cells. 20 tomograms used for comprehensive annotations were acquired with a dose-symmetric tilt scheme^6^ employing similar acquisition parameters (**Extended Data Tables 4-6, Online Methods**), with the only difference being imaging with traditional defocus-only (DEF) or, in addition, the usage of a VPP. All RIBO and FAS particles annotated in the two datasets (DEF and VPP) were subjected to subtomogram analysis including CTF correction utilizing CTF fitting and model creation in Warp, and 3D refinements, hierarchical, and focused 3D classifications in RELION. During this structural analysis we made several observations which will be discussed in the following.

As described previously^7–9^, acquiring tomography data with a VPP emphasizes low frequencies. Pronounced low frequencies lead to improved image contrast which can facilitate data mining. With our method, more particles were detected in VPP than in DEF ground truth annotations of FAS complexes, which represent more challenging targets for data mining due to their low abundance and hollow structural signature. The increased SNR in VPP was also advantageous in hierarchical classifications of ribosomes (**Supplementary Fig. 2**), while no difference was apparent for FAS likely due to the low particle numbers (**Supplementary Fig. 3**). For VPP, all ribosomes clustered in defined averages which confirmed the high performance of DeePiCt and enabled the classification of a subset of 60S large subunits (**Supplementary Fig. 2**). In comparison, most DEF ribosomes from ground truth annotations or DeePiCt predictions clustered in poorly defined classes and no 60S large subunits could be separated even when classification results were improved by optimized particle poses (coordinates and orientations) after multi-particle refinement in M^10^ **(Supplementary Fig. 4, 5**). Also, in classifications focused on an area close to the head of the ribosomal small subunit and at the peptide exit tunnel the additional head density was already apparent in 2D slices of the respective classes in the VPP data (**Supplementary Fig. 6 and 7**, respectively). Similarly, ribosomes close to ER and mitochondria that clustered into classes with adjacent membrane densities are more pronounced in VPP (**Supplementary Fig. 8 and 9**, respectively).

However, fine structural details that provide insights into functional configurations of molecular complexes are lost or cannot be recovered in the final reconstructions of VPP tomography data^9^. This has been suggested to be caused by inaccurate weighing^9^ and a signal loss at high spatial frequencies^11^. The preservation of high-resolution information in DEF in comparison to VPP becomes especially apparent when looking at fine structural details in the subtomogram averages. Densities fitting ACPs of FAS (**Extended data Fig. 8 d-e**), P-site tRNAs (**Extended data Fig. 8 j-k**) inside the 80S ribosome and densities connecting ribosomes to an adjacent mitochondrion membrane (**Supplementary Fig. 9**) could only be resolved in DEF data.

We conclude that the usage of a VPP in tomography acquisition facilitated data mining including the demonstrated domain generalization of applying DeePiCt models trained in VPP to DEF, and improved 3D classifications. This makes it a valuable tool that ultimately provides benefits at the exploratory phase of structural analysis, helping the design of experiments and analysis pipelines that can eventually be performed on DEF data from which higher resolution reconstructions can be obtained.

---

**Supplementary Fig. 1.**
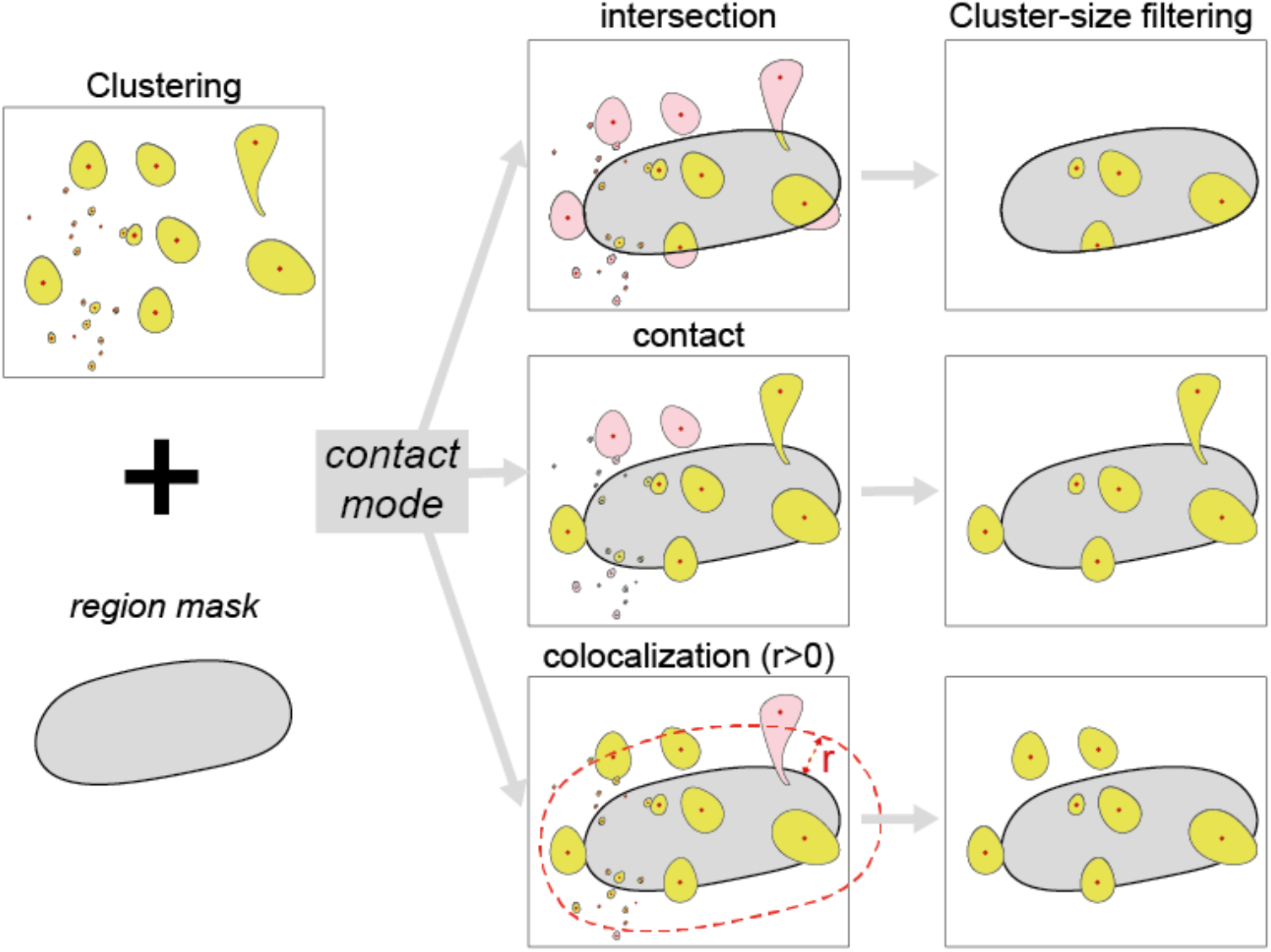
3D CNN post-processing steps. The clustered 3D CNN output (yellow) and a *region mask* (white) are combined according to the three contact modes. Cluster centroids are represented by the red dots.

**Supplementary Fig. 2.**
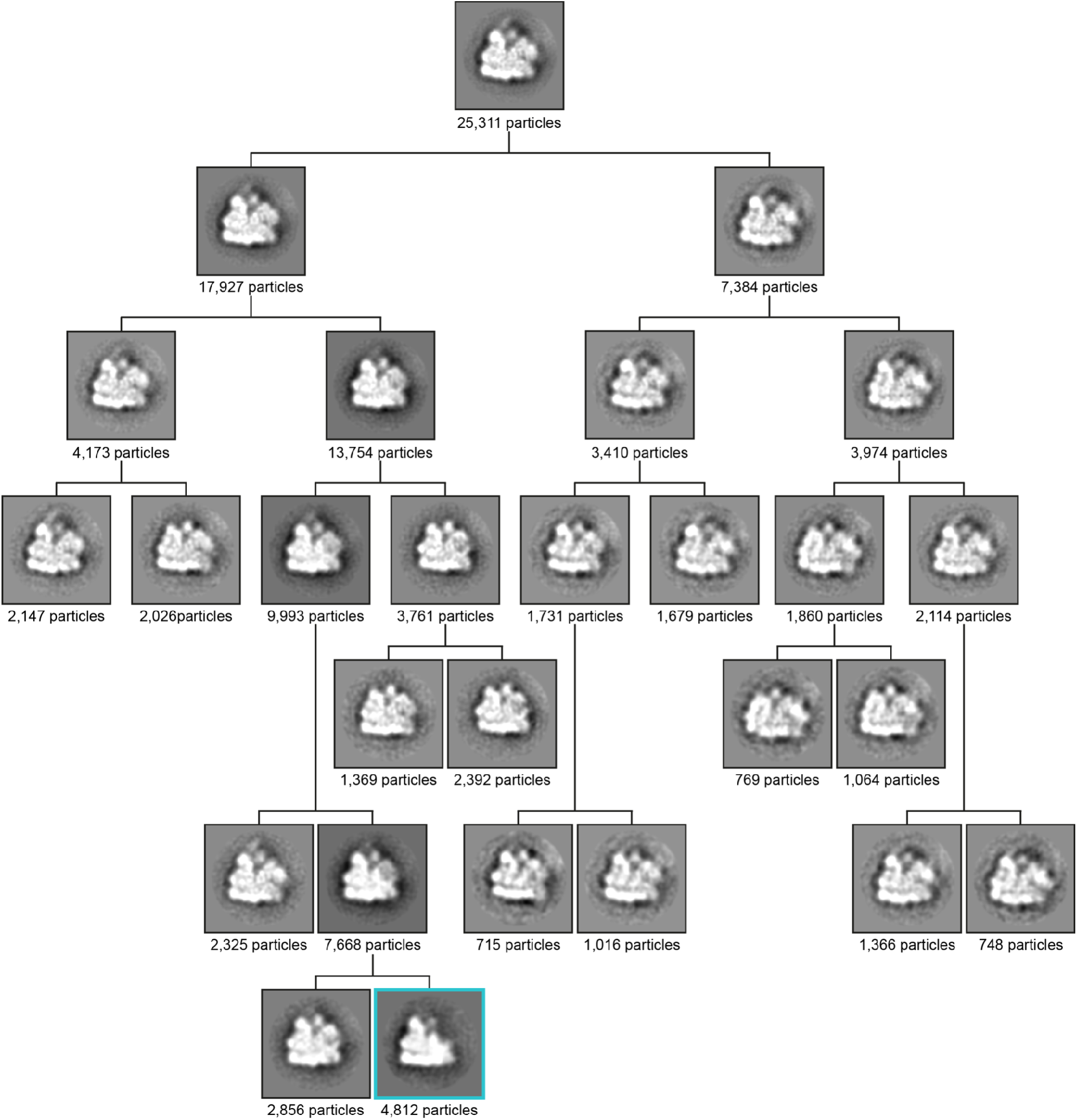
Hierarchical 3D classification of cytosolic ribosomes in VPP ground truth. 2D slices through 3D class averages of iteration 25 are displayed. The highlighted class (cyan box) shows the 60S large ribosomal subunit class.

**Supplementary Fig. 3.**
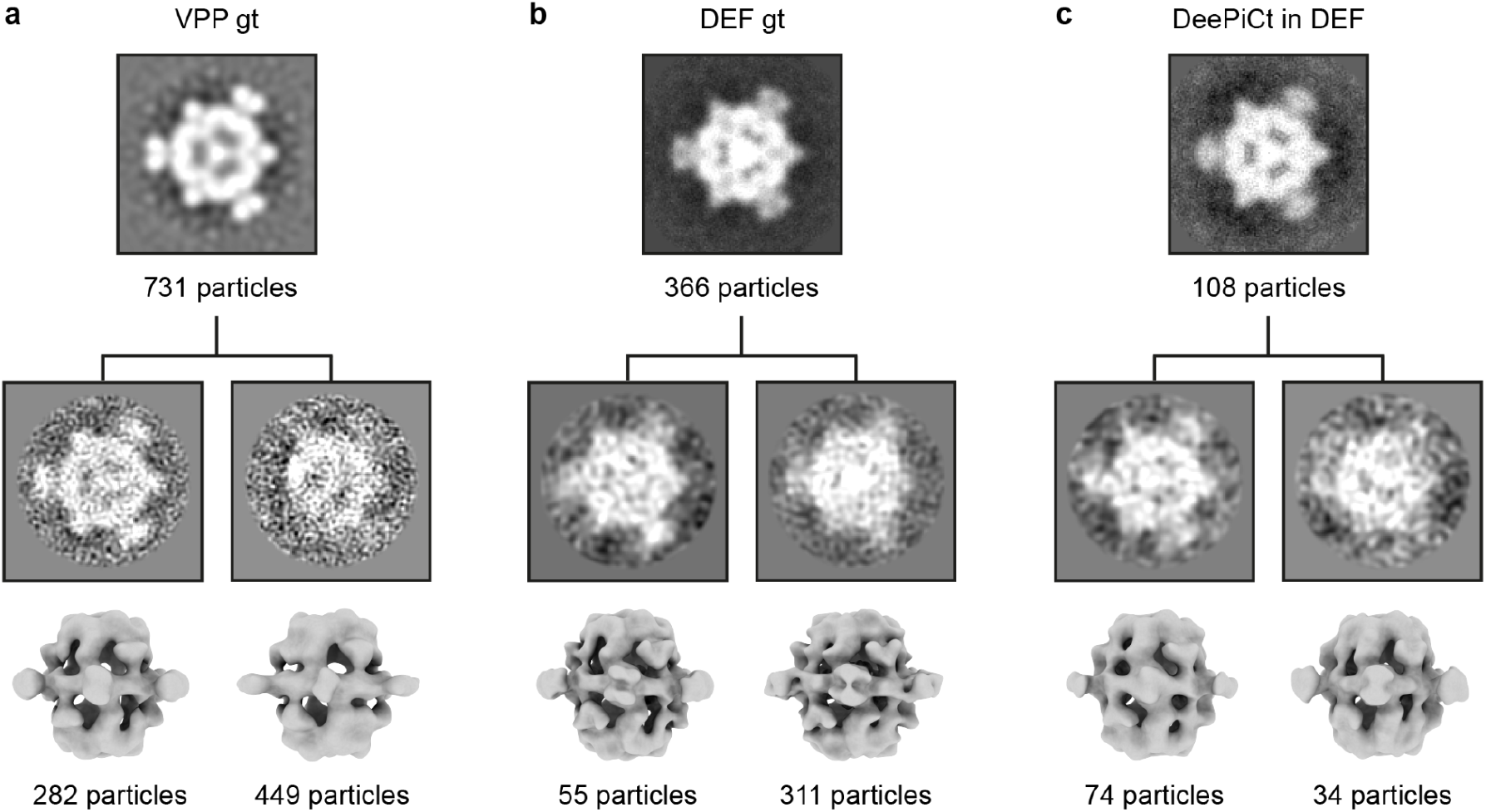
3D classification of VPP and DEF ground truth, and DeePiCt prediction in DEF dataset for FAS. **a-c.** 2D slices through 3D-refined subtomogram average of FAS using D3 symmetry recovered from VPP ground truth (gt), DEF gt or DeePiCt prediction on DEF datasets. 3D classifications separated in all 3 datasets into two classes of which one is better structurally defined than the other. 3D refinements for DEF were performed in M.

**Supplementary Fig. 4.**
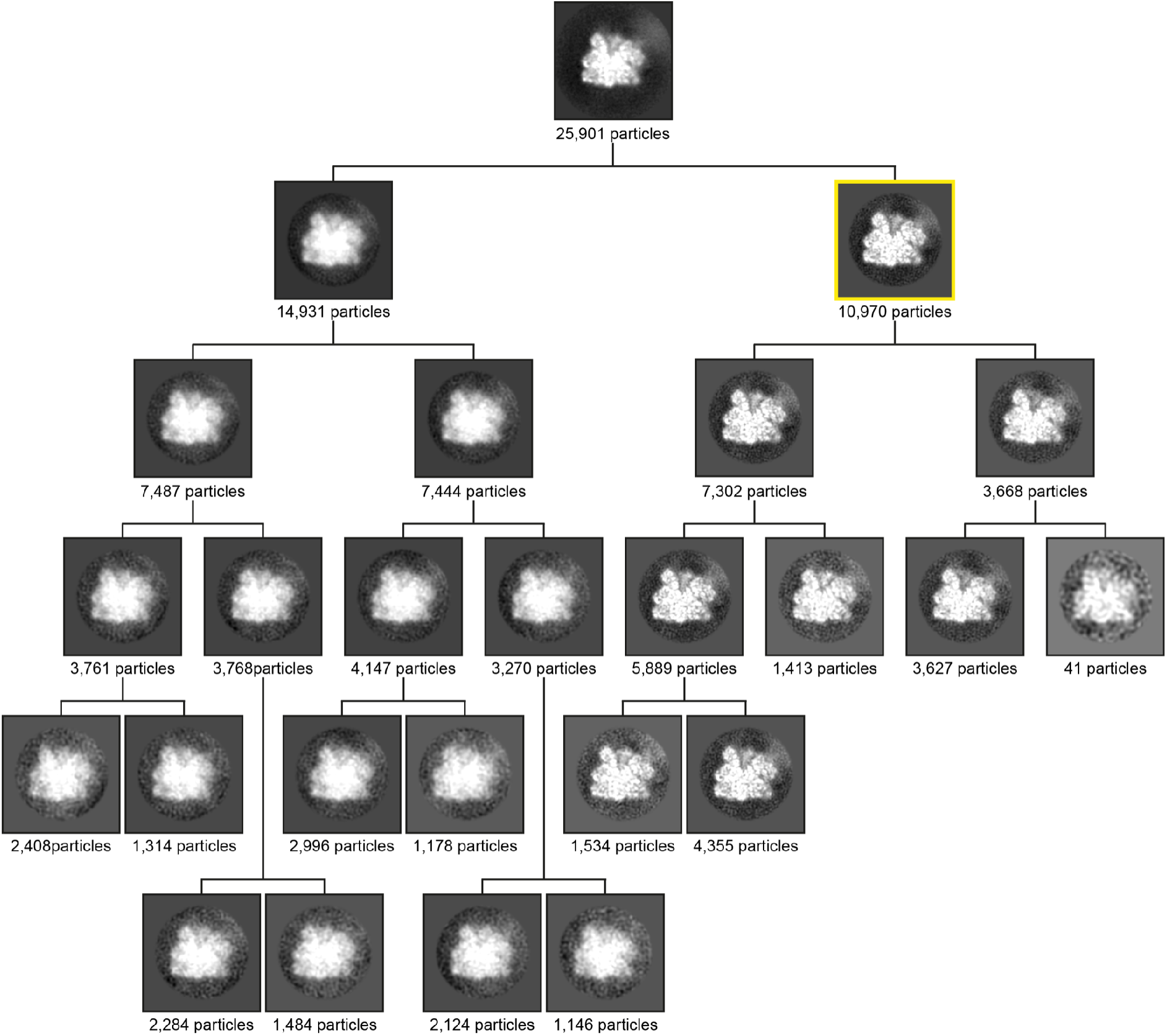
Hierarchical 3D classification of cytosolic ribosomes in DEF ground truth starting from M-refined alignments. 2D slices through 3D class averages of iteration 25 are displayed. The highlighted class (yellow box) was further refined in M, resulting in a well aligned class with a resolution of 9.3 Å.

**Supplementary Fig. 5.**
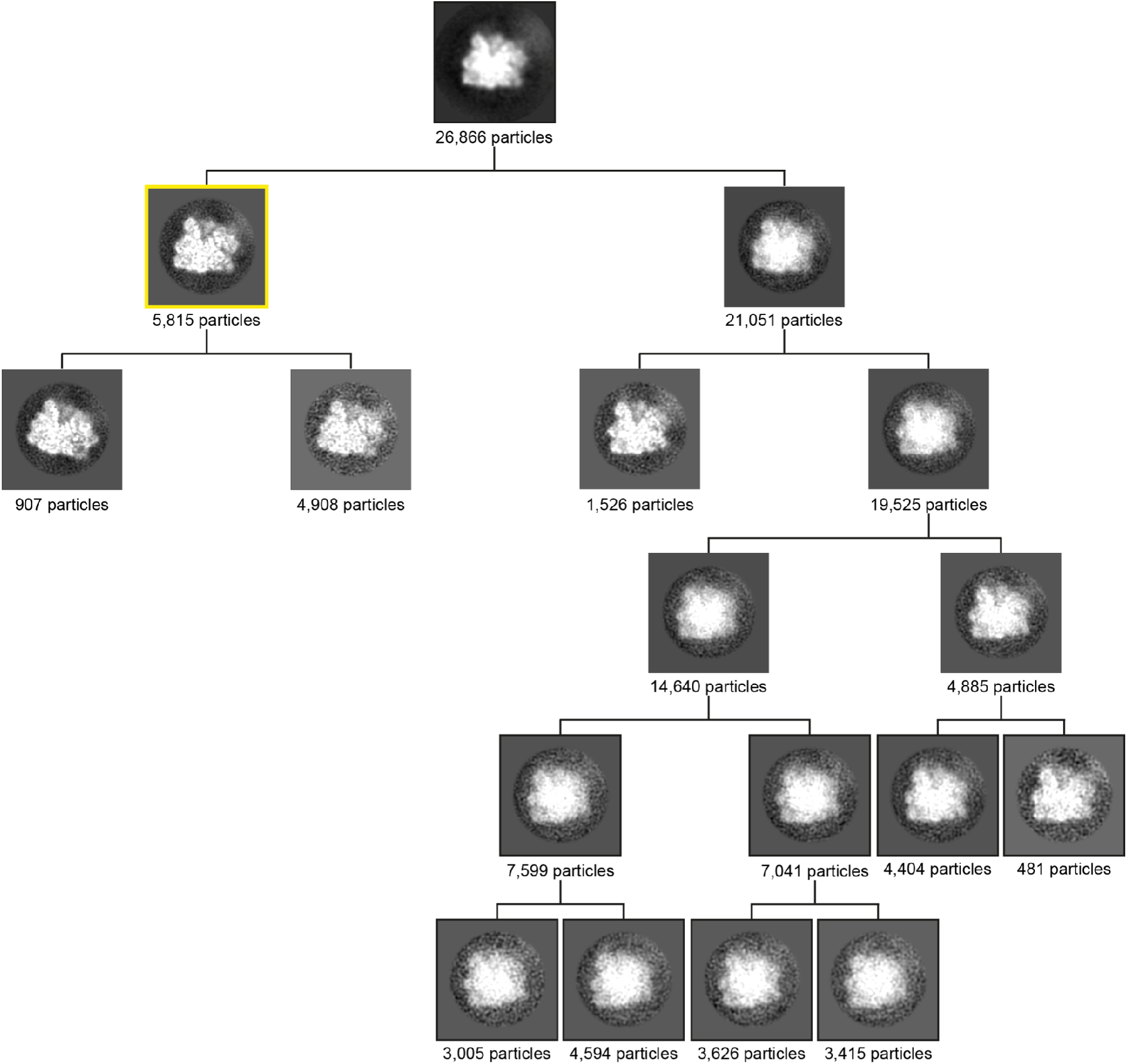
Hierarchical 3D classification of cytosolic ribosomes from DeePiCt predictions in DEF data starting from M-refined alignments. 2D slices through 3D class averages of iteration 25 are displayed. The highlighted class (yellow box) was further refined in M resulting in a well-aligned class with a resolution of 9.4 Å. The right side of the classification representing the vast majority of ribosomes does not converge into a well defined map.

**Supplementary Fig. 6.**
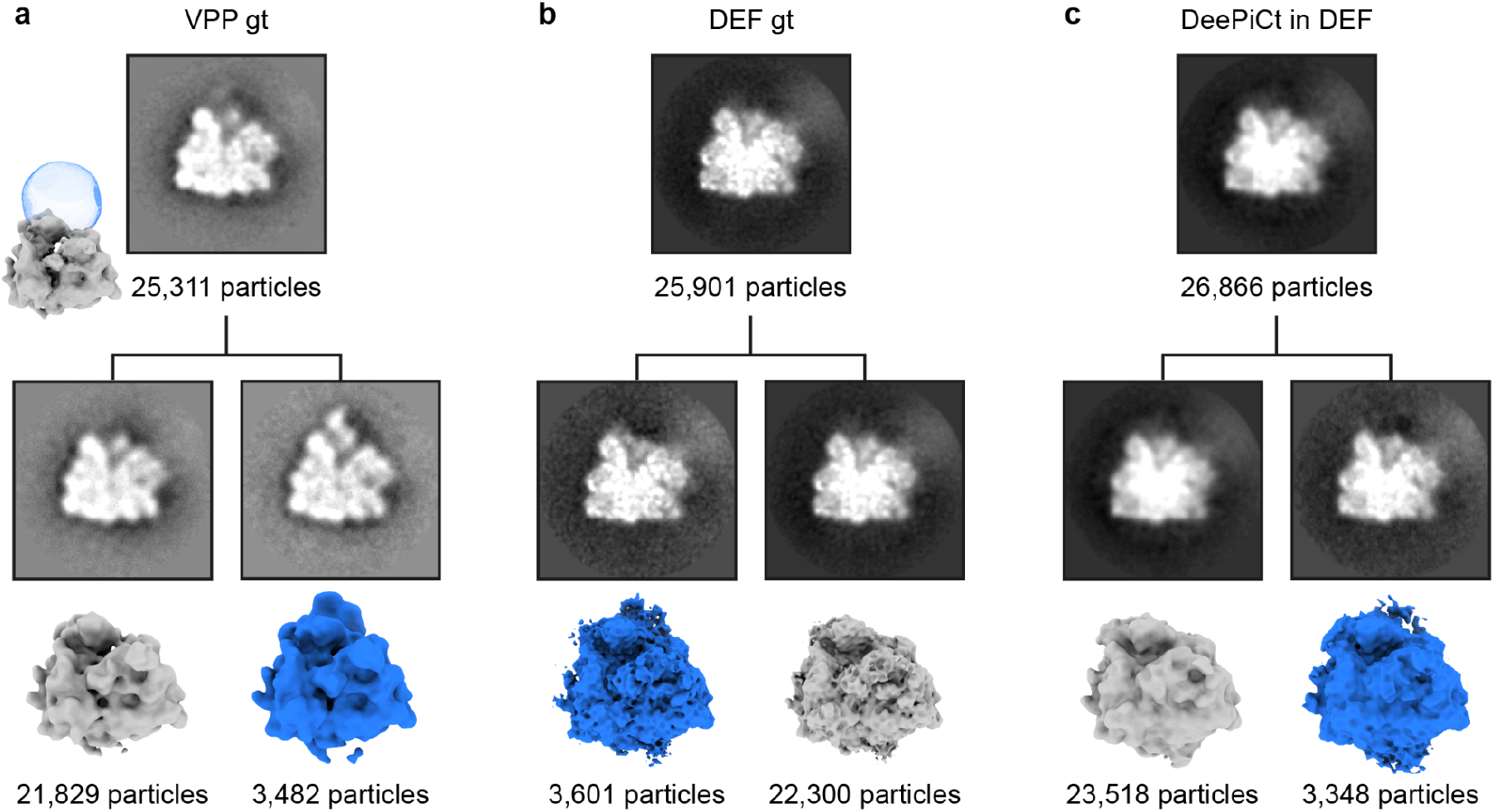
3D classifications and refinements of cryo-EM densities in proximity to the head of the 40S small ribosomal subunit in VPP gt, DEF gt, and DeePiCt predictions in DEF datasets. **a-c.** 2D slices through 3D-refined subtomogram average of all cytosolic ribosomes from VPP ground truth (gt), DEF gt or DeePiCt predictions on DEF datasets. Focused 3D classifications (blue sphere mask displayed in a) separated in all 3 datasets one class (dark blue) with an additional density close to the head of the 40S small ribosomal subunit. 3D refinements for DEF were performed in M.

**Supplementary Fig. 7.**
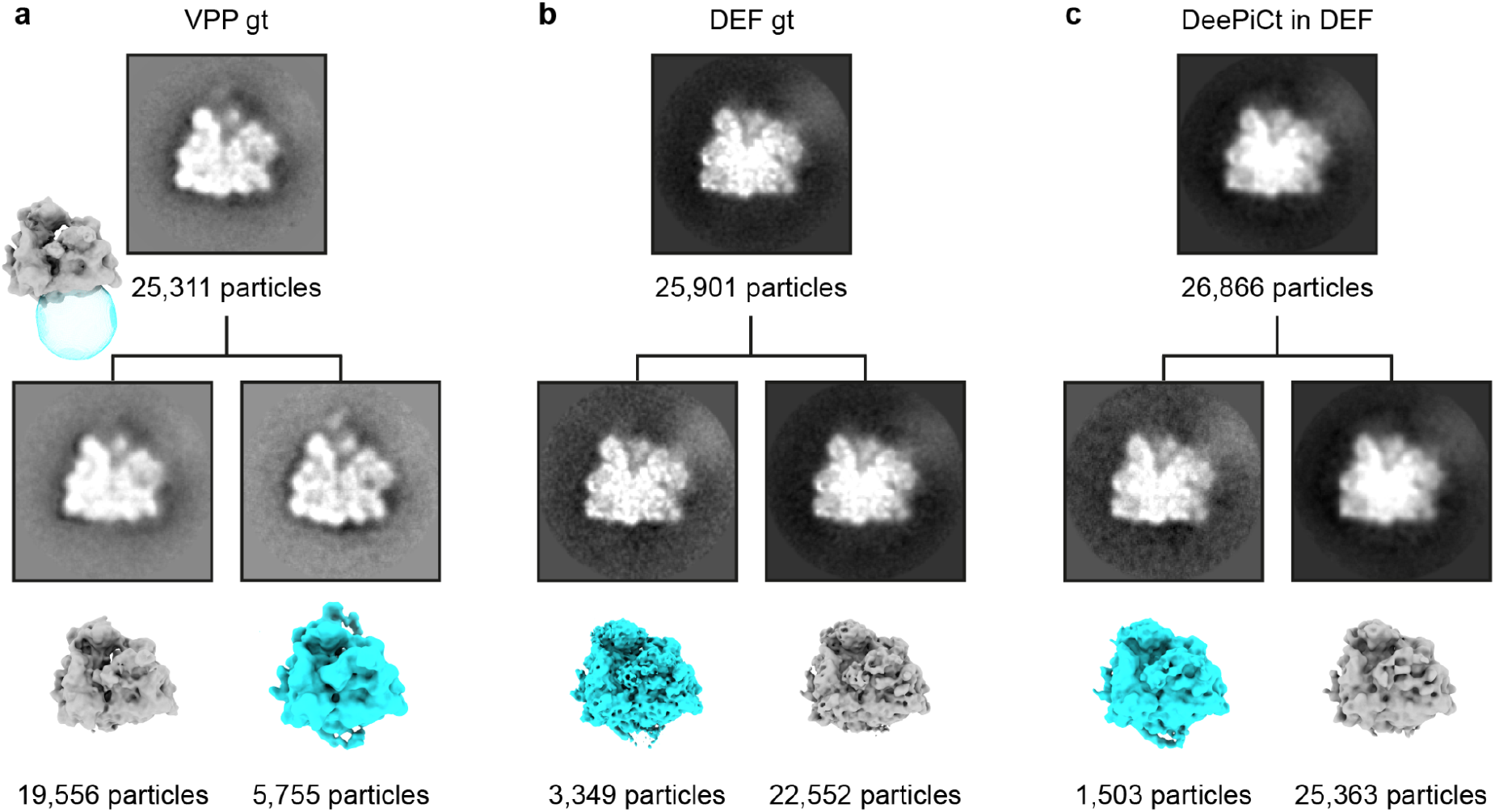
3D classifications and refinements of ribosomes at the exit tunnel in VPP gt, DEF gt, and DeePiCt predictions in DEF datasets. **a-c.** 2D slices through 3D-refined subtomogram average of all ribosomes from VPP ground truth (gt), DEF gt or DeePiCt prediction on DEF datasets. Focused 3D classifications (cyan sphere mask displayed in a) separated in all 3 datasets one class (cyan) with an additional exit tunnel density. 3D refinements for DEF were performed in M.

**Supplementary Fig. 8.**
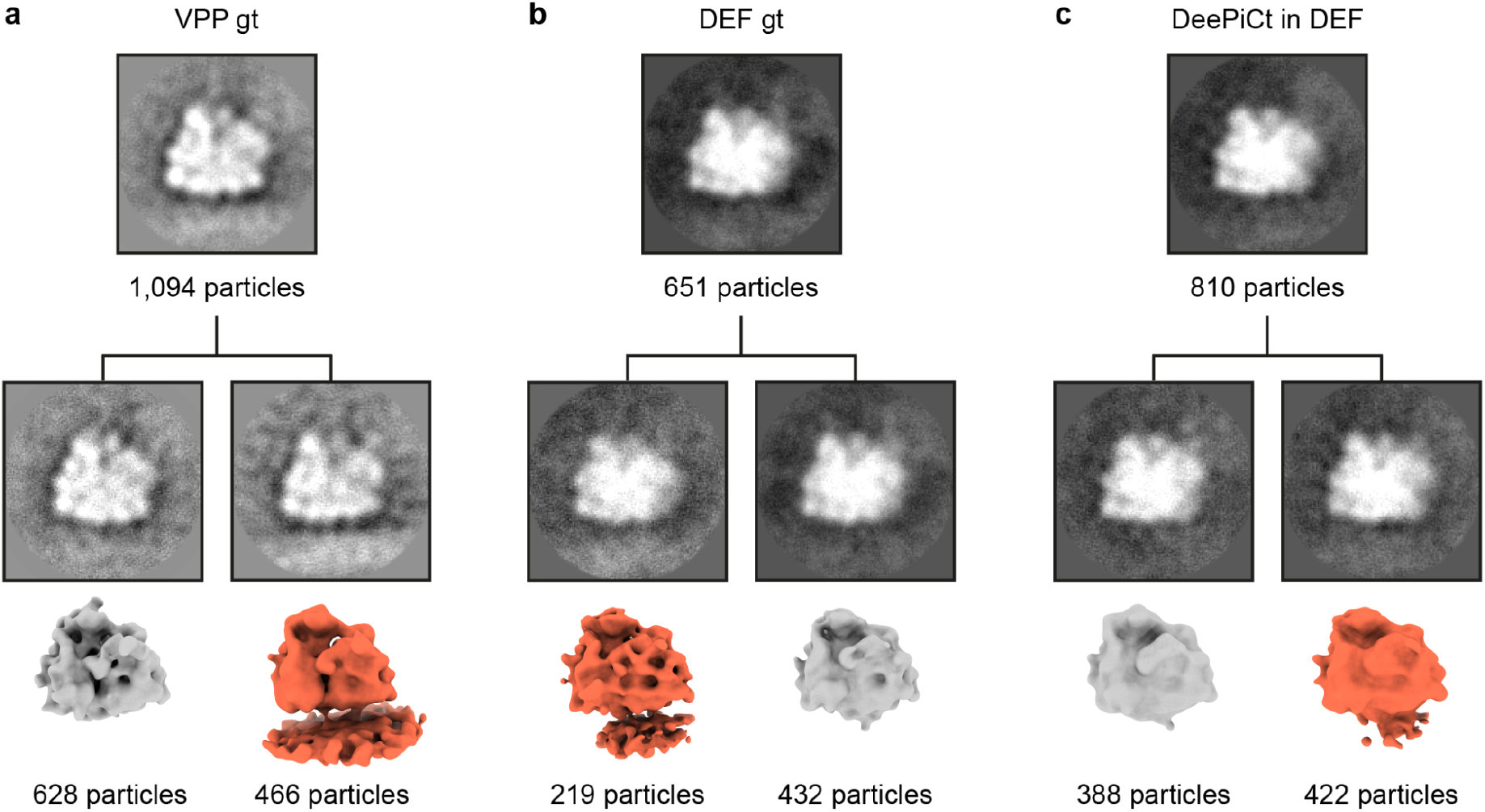
3D classifications and refinements of ER-bound ribosomes in VPP gt, DEF gt, and DeePiCt predictions in DEF datasets. **a-c.** 2D slices through 3D-refined subtomogram average of ribosomes in 25 nm distance to the ER recovered from VPP ground truth (gt), DEF gt or DeePiCt predictions on DEF datasets. Focused 3D classifications (cyan sphere mask displayed in Supplementary Fig. 7a) separated in all 3 datasets one class (orange) with membrane density. 3D refinements for DEF were performed in M.

**Supplementary Fig. 9.**
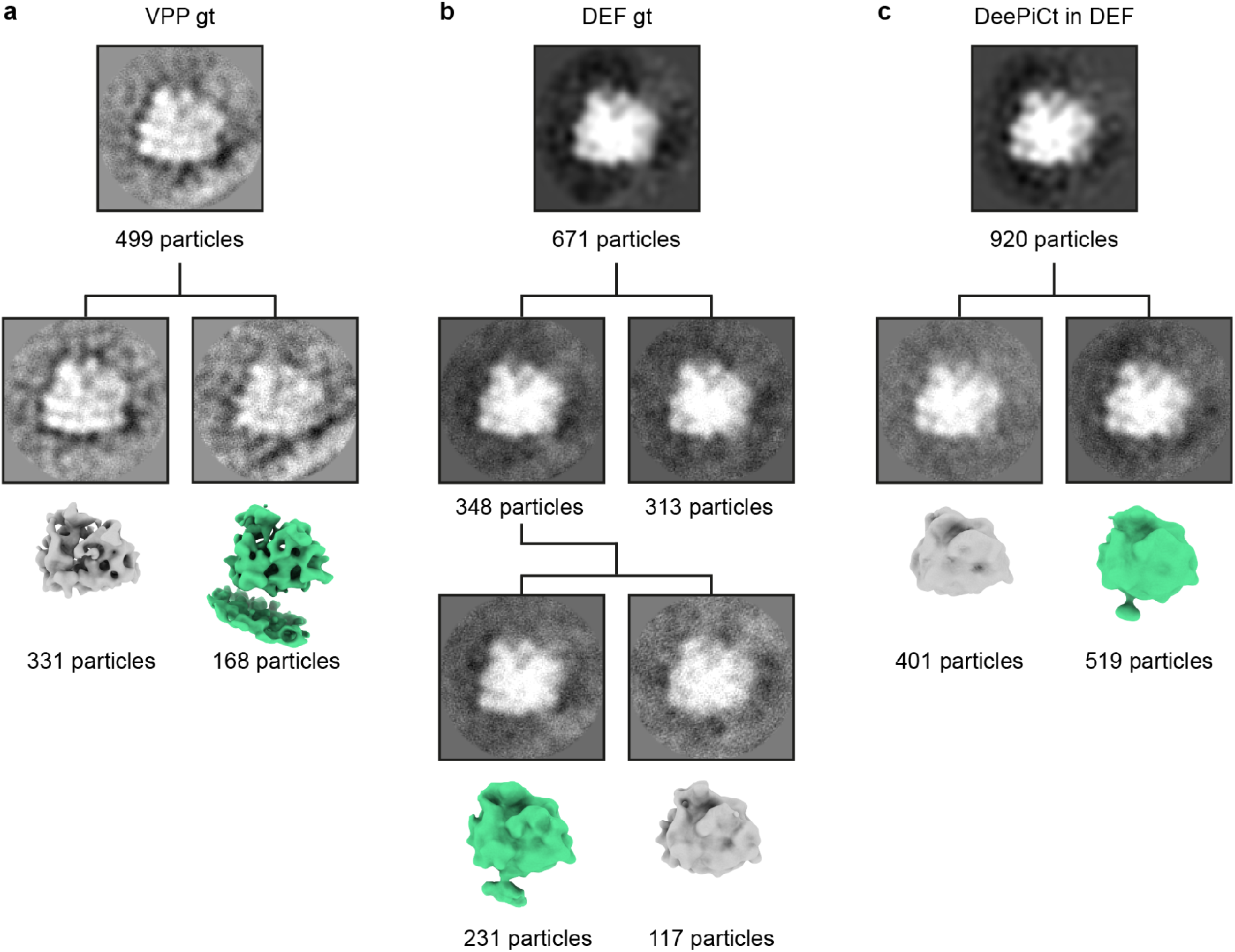
3D classifications and refinements of mitochondria-bound ribosomes in VPP gt, DEF gt, and DeePiCt predictions in DEF datasets. **a-c.** 2D slices through 3D-refined subtomogram average of ribosomes in 25 nm distance to the mitochondria recovered from VPP ground truth (gt), DEF gt or DeePiCt prediction on DEF datasets. Focused 3D classifications (cyan sphere mask displayed in Supplementary Fig. 7a) separated in all 3 datasets one class (green) with membrane density. 3D refinements for DEF were performed in M.

## Bibliography

1. Mahamid, J. et al. Visualizing the molecular sociology at the HeLa cell nuclear periphery. Science 351, 969–972 (2016).

2. Pfeffer, S. et al. Dissecting the molecular organization of the translocon-associated protein complex. Nat. Commun. 8, 14516 (2017).

3. Albert, S. et al. Proteasomes tether to two distinct sites at the nuclear pore complex. Proc Natl Acad Sci USA 114, 13726–13731 (2017).

4. Bäuerlein, F. J. B. et al. In situ architecture and cellular interactions of polyq inclusions. Cell 171, 179–187.e10 (2017).

5. Guo, Q. et al. In Situ Structure of Neuronal C9orf72 Poly-GA Aggregates Reveals Proteasome Recruitment. Cell 172, 696–705.e12 (2018).

6. Albert, S. et al. Direct visualization of degradation microcompartments at the ER membrane. Proc Natl Acad Sci USA 117, 1069–1080 (2020).

7. Wilfling, F. et al. A Selective Autophagy Pathway for Phase-Separated Endocytic Protein Deposits. Mol. Cell 80, 764–778.e7 (2020).

8. Watanabe, R. et al. The In Situ Structure of Parkinson’s Disease-Linked LRRK2. Cell 182, 1508–1518.e16 (2020).

9. Böhning, J. & Bharat, T. A. M. Towards high-throughput in situ structural biology using electron cryotomography. Prog. Biophys. Mol. Biol. 160, 97–103 (2021).

10. Weis, F. & Hagen, W. J. H. Combining high throughput and high quality for cryo-electron microscopy data collection. Acta Crystallogr. D Struct. Biol. 76, 724–728 (2020).

11. Klumpe, S. et al. A modular platform for automated cryo-FIB workflows. eLife 10, (2021).

12. Zachs, T. et al. Fully automated, sequential focused ion beam milling for cryo-electron tomography. eLife 9, (2020).

13. Pyle, E. & Zanetti, G. Current data processing strategies for cryo-electron tomography and subtomogram averaging. Biochem. J. 478, 1827–1845 (2021).

14. Lučič, V., Rigort, A. & Baumeister, W. Cryo-electron tomography: the challenge of doing structural biology in situ. J. Cell Biol. 202, 407–419 (2013).

15. Wan, W. & Briggs, J. A. G. Cryo-Electron Tomography and Subtomogram Averaging. Meth. Enzymol. 579, 329–367 (2016).

16. Wu, X., Zeng, X., Zhu, Z., Gao, X. & Xu, M. Template-Based and Template-Free Approaches in Cellular Cryo-Electron Tomography Structural Pattern Mining. in Computational Biology (ed. Husi, H.) (Codon Publications, 2019). doi:10.15586/computationalbiology.2019.ch11.

17. Bohm, J. et al. Toward detecting and identifying macromolecules in a cellular context: template matching applied to electron tomograms. Proc Natl Acad Sci USA 97, 14245–14250 (2000).

18. Bäuerlein, F. J. B. & Baumeister, W. Towards visual proteomics at high resolution. J. Mol. Biol. 433, 167187 (2021).

19. Xu, M., Beck, M. & Alber, F. Template-free detection of macromolecular complexes in cryo electron tomograms. Bioinformatics 27, i69–76 (2011).

20. Martinez-Sanchez, A. et al. Template-free detection and classification of membrane-bound complexes in cryo-electron tomograms. Nat. Methods 17, 209–216 (2020).

21. Krizhevsky, A., Sutskever, I. & Hinton, G. E. ImageNet classification with deep convolutional neural networks. Commun. ACM 60, 84–90 (2012).

22. LeCun, Y. Deep learning & convolutional networks. in 2015 IEEE Hot Chips 27 Symposium (HCS) 1–95 (IEEE, 2015). doi:10.1109/HOTCHIPS.2015.7477328.

23. LeCun, Y., Bengio, Y. & Hinton, G. Deep learning. Nature 521, 436–444 (2015).

24. Chen, M. et al. Convolutional neural networks for automated annotation of cellular cryo-electron tomograms. Nat. Methods 14, 983–985 (2017).

25. Ronneberger, O., Fischer, P. & Brox, T. U-Net: Convolutional Networks for Biomedical Image Segmentation. in Medical Image Computing and Computer-Assisted Intervention (MICCAI) (eds. Navab, N., Hornegger, J., Wells, W. M. & Frangi, A. F.) vol. 9351 234–241 (Springer International Publishing, 2015).

26. Gubbins, I. et al. Classification in Cryo-Electron Tomograms. https://diglib.eg.org/handle/10.2312/3dor20191061.

27. Moebel, E., Martinez, A., Larivière, D. & Ortiz, J. 3D ConvNet improves macromolecule localization in 3D cellular cryo-electron tomograms. (2018).

28. Moebel, E. et al. Deep learning improves macromolecule identification in 3D cellular cryo-electron tomograms. Nat. Methods 18, 1386–1394 (2021).

29. Gubins, I. et al. SHREC 2021: Classification in Cryo-electron Tomograms. The Eurographics Association (2021) doi:10.2312/3dor.20211307.

30. Srivastava, N., Hinton, G., Krizhevsky, A., Sutskever, I. & Salakhutdinov, R. Dropout: A Simple Way to Prevent Neural Networks from Overfitting. J Mach Learn Res 15, (2014).

31. . Ioffe, S. Batch Normalization: Accelerating Deep Network Training by Reducing Internal Covariate Shift. in vol. 37 (Association for Computing Machinery, 2015).

32. Tegunov, D. & Cramer, P. Real-time cryo-electron microscopy data preprocessing with Warp. Nat. Methods 16, 1146–1152 (2019).

33. Tegunov, D., Xue, L., Dienemann, C., Cramer, P. & Mahamid, J. Multi-particle cryo-EM refinement with M visualizes ribosome-antibiotic complex at 3.5 Å in cells. Nat. Methods 18, 186–193 (2021).

34. Zivanov, J. et al. New tools for automated high-resolution cryo-EM structure determination in RELION-3. eLife 7, (2018).

35. Castaño-Díez, D. The Dynamo package for tomography and subtomogram averaging: components for MATLAB, GPU computing and EC2 Amazon Web Services. Acta Crystallogr. D Struct. Biol. 73, 478–487 (2017).

36. Hrabe, T. et al. PyTom: a python-based toolbox for localization of macromolecules in cryo-electron tomograms and subtomogram analysis. J. Struct. Biol. 178, 177–188 (2012).

37. . Chen, M. et al. A complete data processing workflow for cryo-ET and subtomogram averaging. Nat. Methods 16, 1161–1168 (2019).

38. Danev, R., Buijsse, B., Khoshouei, M., Plitzko, J. M. & Baumeister, W. Volta potential phase plate for in-focus phase contrast transmission electron microscopy. Proc Natl Acad Sci USA 111, 15635– 15640 (2014).

39. Danev, R., Tegunov, D. & Baumeister, W. Using the Volta phase plate with defocus for cryo-EM single particle analysis. eLife 6, (2017).

40. Zimmerli, C. E. et al. Nuclear pores dilate and constrict in cellulo. Science 374, eabd9776 (2021).

41. Carpy, A. et al. Absolute proteome and phosphoproteome dynamics during the cell cycle of Schizosaccharomyces pombe (Fission Yeast). Mol. Cell. Proteomics 13, 1925–1936 (2014).

42. Marguerat, S. et al. Quantitative analysis of fission yeast transcriptomes and proteomes in proliferating and quiescent cells. Cell 151, 671–683 (2012).

43. Turoňová, B. et al. Benchmarking tomographic acquisition schemes for high-resolution structural biology. Nat. Commun. 11, 876 (2020).

44. Leibundgut, M., Jenni, S., Frick, C. & Ban, N. Structural basis for substrate delivery by acyl carrier protein in the yeast fatty acid synthase. Science 316, 288–290 (2007).

45. Snowden, J. S. et al. Structural insight into Pichia pastoris fatty acid synthase. Sci. Rep. 11, 9773 (2021).

46. Gipson, P. et al. Direct structural insight into the substrate-shuttling mechanism of yeast fatty acid synthase by electron cryomicroscopy. Proc Natl Acad Sci USA 107, 9164–9169 (2010).

47. Singh, K. et al. Discovery of a regulatory subunit of the yeast fatty acid synthase. Cell 180, 1130–1143.e20 (2020).

48. Kastritis, P. L. et al. Capturing protein communities by structural proteomics in a thermophilic eukaryote. Mol. Syst. Biol. 13, 936 (2017).

49. . Jenni, S. et al. Structure of fungal fatty acid synthase and implications for iterative substrate shuttling. Science 316, 254–261 (2007).

50. Ranjan, N. et al. Yeast translation elongation factor eEF3 promotes late stages of tRNA translocation. EMBO J. 40, e106449 (2021).

51. Andersen, C. B. F. et al. Structure of eEF3 and the mechanism of transfer RNA release from the E-site. Nature 443, 663–668 (2006).

52. Becker, T. et al. Structure of monomeric yeast and mammalian Sec61 complexes interacting with the translating ribosome. Science 326, 1369–1373 (2009).

53. Armache, J.-P. et al. Cryo-EM structure and rRNA model of a translating eukaryotic 80S ribosome at 5.5-A resolution. Proc Natl Acad Sci USA 107, 19748–19753 (2010).

54. Shankar, V. et al. rRNA expansion segment 27Lb modulates the factor recruitment capacity of the yeast ribosome and shapes the proteome. Nucleic Acids Res. 48, 3244–3256 (2020).

55. Fujii, K., Susanto, T. T., Saurabh, S. & Barna, M. Decoding the function of expansion segments in ribosomes. Mol. Cell 72, 1013–1020.e6 (2018).

56. Greber, B. J. et al. Insertion of the Biogenesis Factor Rei1 Probes the Ribosomal Tunnel during 60S Maturation. Cell 164, 91–102 (2016).

57. Wild, K. et al. MetAP-like Ebp1 occupies the human ribosomal tunnel exit and recruits flexible rRNA expansion segments. Nat. Commun. 11, 776 (2020).

58. Kowalinski, E. et al. The crystal structure of Ebp1 reveals a methionine aminopeptidase fold as binding platform for multiple interactions. FEBS Lett. 581, 4450–4454 (2007).

59. Gold, V. A., Chroscicki, P., Bragoszewski, P. & Chacinska, A. Visualization of cytosolic ribosomes on the surface of mitochondria by electron cryo-tomography. EMBO Rep. 18, 1786–1800 (2017).

60. Avendaño-Monsalve, M. C., Ponce-Rojas, J. C. & Funes, S. From cytosol to mitochondria: the beginning of a protein journey. Biol. Chem. 401, 645–661 (2020).

61. Tucker, K. & Park, E. Cryo-EM structure of the mitochondrial protein-import channel TOM complex at near-atomic resolution. Nat. Struct. Mol. Biol. 26, 1158–1166 (2019).

62. Lesnik, C., Cohen, Y., Atir-Lande, A., Schuldiner, M. & Arava, Y. OM14 is a mitochondrial receptor for cytosolic ribosomes that supports co-translational import into mitochondria. Nat. Commun. 5, 5711 (2014).

63. George, R., Walsh, P., Beddoe, T. & Lithgow, T. The nascent polypeptide-associated complex (NAC) promotes interaction of ribosomes with the mitochondrial surface in vivo. FEBS Lett. 516, 213–216 (2002).

64. Stalling, D., Westerhoff, M. & Hege, H.-C. amira: A Highly Interactive System for Visual Data Analysis. in Visualization Handbook 749–767 (Elsevier, 2005). doi:10.1016/B978-012387582-2/50040-X.

65. Vignaud, T. et al. Stress fibres are embedded in a contractile cortical network. Nat. Mater. 20, 410– 420 (2021).

66. Bharat, T. A. M. & Scheres, S. H. W. Resolving macromolecular structures from electron cryo-tomography data using subtomogram averaging in RELION. Nat. Protoc. 11, 2054–2065 (2016).

67. Wagner, F. R. et al. Preparing samples from whole cells using focused-ion-beam milling for cryo-electron tomography. Nat. Protoc. 15, 2041–2070 (2020).

68. Schaffer, M. et al. Optimized cryo-focused ion beam sample preparation aimed at in situ structural studies of membrane proteins. J. Struct. Biol. 197, 73–82 (2017).

69. Mastronarde, D. N. Advanced data acquisition from electron microscopes with serialem. Microsc. Microanal. 24, 864–865 (2018).

70. Hagen, W. J. H., Wan, W. & Briggs, J. A. G. Implementation of a cryo-electron tomography tilt-scheme optimized for high resolution subtomogram averaging. J. Struct. Biol. 197, 191–198 (2017).

71. Allegretti, M. et al. In-cell architecture of the nuclear pore and snapshots of its turnover. Nature 586, 796–800 (2020).

72. Mastronarde, D. N. & Held, S. R. Automated tilt series alignment and tomographic reconstruction in IMOD. J. Struct. Biol. 197, 102–113 (2017).

73. Tang, G. et al. EMAN2: an extensible image processing suite for electron microscopy. J. Struct. Biol. 157, 38–46 (2007).

74. Wu, J.-Q. & Pollard, T. D. Counting cytokinesis proteins globally and locally in fission yeast. Science 310, 310–314 (2005).

75. Mitchison, J. M. The growth of single cells. Exp. Cell Res. 13, 244–262 (1957).

76. Nurse, P. Genetic control of cell size at cell division in yeast. Nature 256, 547–551 (1975).

77. Turonova. turonova/novaSTA: novaSTA. Zenodo (2020) doi:10.5281/zenodo.3973623.

78. Rigort, A. et al. Automated segmentation of electron tomograms for a quantitative description of actin filament networks. J. Struct. Biol. 177, 135–144 (2012).

79. Redemann, S. et al. The segmentation of microtubules in electron tomograms using Amira. Methods Mol. Biol. 1136, 261–278 (2014).

80. Pettersen, E. F. et al. UCSF Chimera—a visualization system for exploratory research and analysis. J. Comput. Chem. 25, 1605–1612 (2004).

81. Goddard, T. D. et al. UCSF ChimeraX: Meeting modern challenges in visualization and analysis.Protein Sci. 27, 14–25 (2018).

82. Moebel, E. et al. Deep Learning Improves Macromolecules Localization and Identification in 3D Cellular Cryo-Electron Tomograms. BioRxiv (2020) doi:10.1101/2020.04.15.042747.

## Bibliography

1. Wu, J.-Q. & Pollard, T. D. Counting cytokinesis proteins globally and locally in fission yeast. Science 310, 310–314 (2005).

2. Mitchison, J. M. The growth of single cells. Exp. Cell Res. 13, 244–262 (1957).

3. Nurse, P. Genetic control of cell size at cell division in yeast. Nature 256, 547–551 (1975).

4. Marguerat, S. et al. Quantitative analysis of fission yeast transcriptomes and proteomes in proliferating and quiescent cells. Cell 151, 671–683 (2012).

5. Carpy, A. et al. Absolute proteome and phosphoproteome dynamics during the cell cycle of Schizosaccharomyces pombe (Fission Yeast). Mol. Cell. Proteomics 13, 1925–1936 (2014).

6. Ranjan, N. et al. Yeast translation elongation factor eEF3 promotes late stages of tRNA translocation. EMBO J. 40, e106449 (2021).

7. Becker, T. et al. Structure of monomeric yeast and mammalian Sec61 complexes interacting with the translating ribosome. Science 326, 1369–1373 (2009).

8. Armache, J.-P. et al. Cryo-EM structure and rRNA model of a translating eukaryotic 80S ribosome at 5.5- A resolution. Proc Natl Acad Sci USA 107, 19748–19753 (2010).

9. Wild, K. et al. MetAP-like Ebp1 occupies the human ribosomal tunnel exit and recruits flexible rRNA expansion segments. Nat. Commun. 11, 776 (2020).

10. Gold, V. A., Chroscicki, P., Bragoszewski, P. & Chacinska, A. Visualization of cytosolic ribosomes on the surface of mitochondria by electron cryo-tomography. EMBO Rep. 18, 1786–1800 (2017).

11. Mahamid, J. et al. Visualizing the molecular sociology at the HeLa cell nuclear periphery. Science 351, 969–972 (2016).

12. Vignaud, T. et al. Stress fibres are embedded in a contractile cortical network. Nat. Mater. 20, 410–420 (2021).

## Bibliography

1. Ronneberger, O., Fischer, P. & Brox, T. U-Net: Convolutional Networks for Biomedical Image Segmentation. in Medical Image Computing and Computer-Assisted Intervention (MICCAI) (eds. Navab, N., Hornegger, J., Wells, W. M. & Frangi, A. F.) vol. 9351 234–241 (Springer International Publishing, 2015).

2. Zeiler, M. D., Krishnan, D., Taylor, G. W. & Fergus, R. Deconvolutional networks. in 2010 IEEE Computer Society Conference on Computer Vision and Pattern Recognition 2528–2535 (IEEE, 2010). doi:10.1109/CVPR.2010.5539957.

3. Milletari, F., Navab, N. & Ahmadi, S.-A. V-Net: Fully Convolutional Neural Networks for Volumetric Medical Image Segmentation. in 2016 Fourth International Conference on 3D Vision (3DV) 565–571 (IEEE, 2016). doi:10.1109/3DV.2016.79.

4. Kingma, D. P. & Ba, J. Adam: A Method for Stochastic Optimization. https://arxiv.org/abs/1412.6980 (2017).

5. van der Walt, S. et al. scikit-image: image processing in Python. PeerJ 2, e453 (2014).

6. Hagen, W. J. H., Wan, W. & Briggs, J. A. G. Implementation of a cryo-electron tomography tilt-scheme optimized for high resolution subtomogram averaging. J. Struct. Biol. 197, 191–198 (2017).

7. Danev, R., Buijsse, B., Khoshouei, M., Plitzko, J. M. & Baumeister, W. Volta potential phase plate for in-focus phase contrast transmission electron microscopy. Proc Natl Acad Sci USA 111, 15635–15640 (2014).

8. Danev, R., Tegunov, D. & Baumeister, W. Using the Volta phase plate with defocus for cryo-EM single particle analysis. eLife 6, (2017).

9. Turoňová, B. et al. Benchmarking tomographic acquisition schemes for high-resolution structural biology. Nat. Commun. 11, 876 (2020).

10. Tegunov, D., Xue, L., Dienemann, C., Cramer, P. & Mahamid, J. Multi-particle cryo-EM refinement with M visualizes ribosome-antibiotic complex at 3.5 Å in cells. Nat. Methods 18, 186–193 (2021).

11. Buijsse, B., Trompenaars, P., Altin, V., Danev, R. & Glaeser, R. M. Spectral DQE of the Volta phase plate. Ultramicroscopy 218, 113079 (2020).

